# In situ structures from relaxed cardiac myofibrils reveal the organization of the muscle thick filament

**DOI:** 10.1101/2023.04.11.536387

**Authors:** Davide Tamborrini, Zhexin Wang, Thorsten Wagner, Sebastian Tacke, Markus Stabrin, Michael Grange, Ay Lin Kho, Martin Rees, Pauline Bennett, Mathias Gautel, Stefan Raunser

**Affiliations:** Department of Structural Biochemistry, Max Planck Institute of Molecular Physiology, 44227 Dortmund, Germany; Randall Centre for Cell and Molecular Biophysics, School of Basic and Medical Biosciences, Kings College London BHF Centre of Research Excellence, New Hunt’s House, Guy’s Campus, London SE1 1UL, UK

## Abstract

The thick filament is a key component of sarcomeres, the basic force-generating and load-bearing unit of striated muscle^1^. Mutations in thick filament proteins are associated with familial hypertrophic cardiomyopathy and other heart and muscle diseases^2, 3^. Despite this central importance for sarcomere force generation, it remains unclear how thick filaments are structurally organized and how its components interact with each other and with thin filaments to enable highly regulated muscle contraction. Here, we present the molecular architecture of native cardiac sarcomeres in the relaxed state, determined by electron cryo-tomography. Our reconstruction of the thick filament reveals the three-dimensional organization of myosin heads and tails, myosin-binding protein C (MyBP-C) and titin, elucidating the structural basis for their interaction during muscle contraction. The arrangement of myosin heads is variable depending on their position along the filament, suggesting that they have different capacities in terms of strain susceptibility and activation. Myosin tails exhibit a distinct arrangement and pattern of interactions. These are likely orchestrated by three alpha and three beta titin chains that are arranged like a spring, suggesting the existence of specialized roles of thick filament segments in length-dependent activation and contraction. Surprisingly, while the three titin alpha chains run along the entire length of the thick filament, titin beta does not. The structure also demonstrates that the C-terminal region of MyBP-C binds myosin tails and unexpectedly also directly interacts with the myosin heads, suggesting a previously undescribed direct role in the preservation of the myosin OFF state. Furthermore, we visualize how MyBP-C forms links between thin and thick filaments. These findings establish a robust groundwork for forthcoming research endeavors aiming to explore muscle disorders that involve sarcomeric structural components.

## Introduction

Muscle contraction requires a shortening of the sarcomere, the basic contractile unit of muscle, caused by the sliding of interdigitating thick and thin filaments. Thick filaments are bipolar structures containing myosin-II, myosin-binding protein C (MyBP-C) and titin, as well as several other proteins near the bare zone, which marks the thick filament’s symmetry axis and the centre of the sarcomere^4^. By convention, this region comprises the P-zone (P stands for proximal), where the so-called crowns of myosin P1, P2 and P3 are located, and the M-band, a bare zone devoid of myosin heads where tails belonging to apposing half-sarcomeres intertwine^5^. Myosin is a 446 kDa complex of a myosin-II dimer whose head or motor domains connect via a neck region to their coiled-coil tails that pack into the thick filament backbone, with each neck bound by an essential and regulatory light chain. Myosin heads are arranged in a quasi-helical array with a 3-fold rotational symmetry in the relaxed state of muscle^6–8^. The pairs of heads of three myosin molecules protrude from the backbone at regular intervals to form a ℌcrown” of heads, with successive crowns axially separated by about 143 Å. The interaction of the myosin motor domains with the thin filament is responsible for force generation and muscle contraction.

MyBP-C localizes in the C-zone which also contains all the myosin motors that are responsible for providing the peak force during contraction^9^. The cardiac isoform of MyBP-C (cMyBP-C) is a 140 kDa protein with 3 fibronectin type 3 (Fn3)-like domains and 8 immunoglobulin (Ig)-like domains that can form links between the thick and thin filaments^10^. Biochemical investigations support a model in which the N-terminal region can bind both actin and myosin heads in a phosphorylation-dependent manner^11, 12^, while the C-terminal region docks to the thick filament via interactions with titin and myosin tails^13, 14^. MyBP-C links can act as mechanical sensor for muscle tension and have both activating and inhibitory effects on the myosin motors, fine tuning the strength and the kinetics of the muscle contraction^15–18^.

Titin is an up to 4.25 MDa protein consisting of up to 169 Ig-like and 132 Fn3-like domains that form a chain from the M-band to the Z-disc, where the actin filaments of opposite polarity are cross-linked by α-actinin, marking the lateral borders of the sarcomeres^19, 20^. Titin acts as a molecular spring that prevents the overstretching of the sarcomere, recoils the sarcomere after stretch release^21, 22^ and is proposed to act as a molecular ruler for assembly of myosin in the A-band (the region of the sarcomere where thin and thick filaments overlap)^23–25^. The exact stoichiometry of titin within the A-band has remained a mystery, with estimates ranging from 2 to 10 molecules per thick filament^26^. With numerous post-translational modification sites scattered along its length and the predicted interactions with myosin and MyBP-C, titin is also proposed to mediate regulatory signals from the cell to the myosin motors^27^.

Models exist for the general arrangement of myosin heads within the cardiac thick filament^28^ and using components reconstituted in vitro, remarkable insights have been gleaned into the regulation of their kinetics^29^. However, how these observations map onto the structure of a fully complemented, natively organised sarcomere, and the molecular interactions that occur within the thick filament in its native environment is still unknown. In particular, knowledge of key molecular players implicated in disease – myosin, titin, MyBP-C – is completely lacking, demonstrating a key gap in understanding of how they carry out their highly regulated functions within the confined space of the thick filament. Here, we present the *in situ* structure of the thick filament in the M-band, P-zone and C-zone in relaxed mouse cardiac myofibrils determined using our established workflow for focused ion beam (FIB) milling and cryo-electron tomography (cryo-ET)^20, 30, 31^. The structure reveals in unprecedented detail and completeness the architecture of the thick filament, visualizing critical interactions between myosin, titin, and MyBP-C, as well as MyBP-C links between thin and thick filaments.

## Results

### Sarcomere structure of relaxed cardiac myofibrils

To obtain a near-native sample suitable for FIB milling and cryo-ET analysis, we relaxed de-membranated mouse cardiac myofibrils in the presence of EGTA, dextran, and mavacamten (see Methods). Mavacamten, an FDA-approved drug to treat hypertrophic cardiomyopathy, stabilizes the OFF (relaxed) state of myosin^32, 33^.

The tomograms showed that the thick filaments were indeed in the OFF state, as defined by the absence of myosin binding to thin filaments (Fig. 1a). Importantly, the tomograms contain structures that could not be visualized *in situ* before. In particular, using sub-tomogram averaging, we could determine the structure of the cardiac thin filament with tropomyosin in the blocked state (myosin binding site occluded) at 8.2 Å resolution (Fig. 1b,d, Extended Data Figs. 1a,b, 2a,b, 3,a,b, Methods) and the structure of the relaxed thick filament (Fig. 1b,c,e, Extended Data Fig. 1,b 2,b Methods). To tease out greater detail in the analysis of the thick filament, we subdivided the filament into overlapping segments and determined their structures independently, obtaining subtle structural differences between the different segments. The resulting reconstructions ranged in resolution from 19.3 Å to 23.6 Å (Extended Data Fig. 2c-h).

**Fig. 1.**
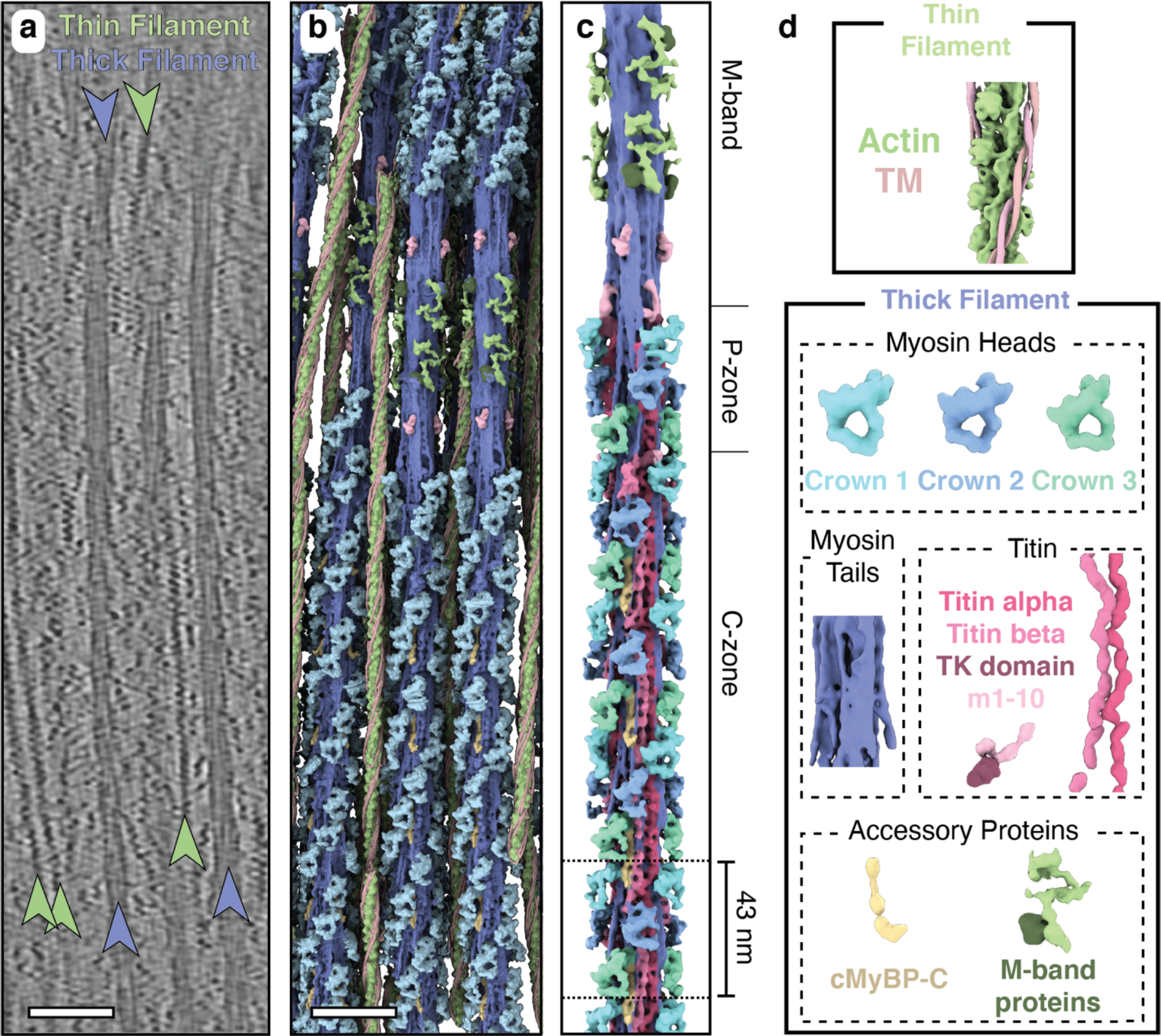
Thick filament structure in the relaxed cardiac sarcomere. **a**, Tomographic slice of a cardiac sarcomere M-band depicting thick and thin filaments. Scale bar, 50 nm. **b**, Reconstructed thick and thin filaments mapped into a tomogram. Thin filaments obstructing the view on the thick filament were removed for clarity. Scale bar, 50 nm **c**, Structure of the thick filament from the M-band to the C-zone. For clarity, only the first four cMyBP-C stripes are shown here. **d**, Illustration of the various sarcomere components and their color code which is maintained throughout the manuscript, unless otherwise indicated. TM, tropomyosin. TK domain, titin kinase domain.

Mapping of the determined structures into their determined XYZ positions within the reconstructed tomograms further enabled us to characterize the 3D arrangement of thin and thick filaments (Fig. 1b, Supplementary Video 1). In agreement with previous structural studies of vertebrate heart muscles, the myosin heads in our combined reconstruction are arranged in a quasi-helical array with a 3-fold rotational symmetry^6–8^ (Fig. 1c, Extended Data Fig. 4). The combined reconstructions comprise the M-band, P-zone, and C-zone and reveal the position of M-band proteins together with the first 31 myosin layers, encompassing 6 titin molecules and 9 cMyBP-C stripes (27 molecules of cMyBP-C) (Fig. 1b,c, 2). This structure of the thick filament has a diameter of 64 nm, a length of 510 nm, and a molecular weight of ∼ 67 MDa and enables a description of the thick filament across the entire sarcomere in a region-by-region specific manner.

### Structural organization of myosin, cMyBP-C and titin

The segments containing the cMyBP-C stripes from 4 to 9 and the titin C-type super-repeats 3-8 have similar structures and were therefore averaged yielding an improved resolution of 18 Å (Fig. 3a, Extended Data Fig. 2h, Methods). We then fitted atomic models of thick filament proteins (obtained either from experimental structures or AlphaFold2 predictions^34, 35^) in the resulting reconstruction, allowing us to assign all densities to the corresponding thick filament components (Fig 3b, Supplementary Video 2).

**Fig. 2.**
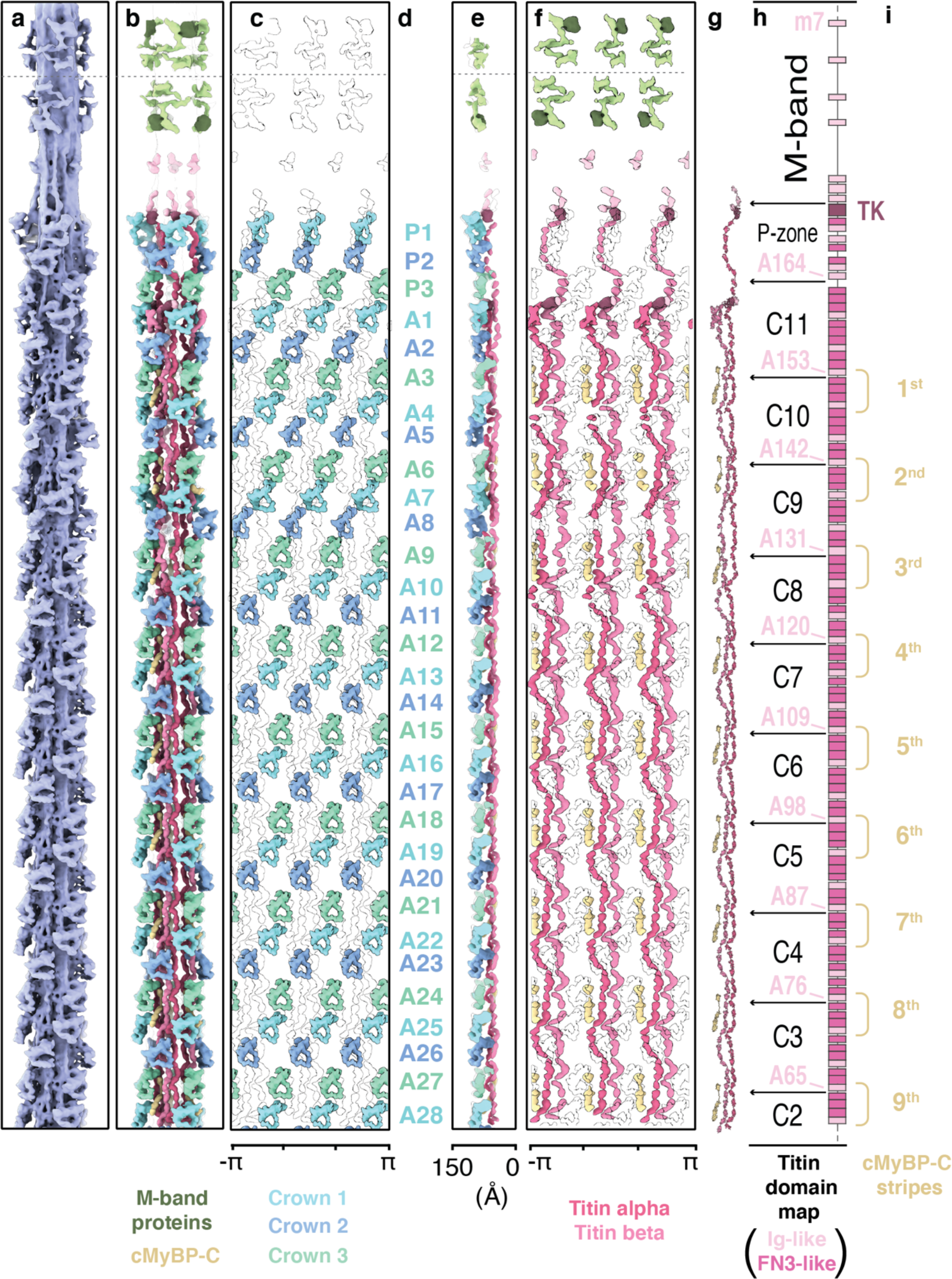
Arrangement of myosin heads, cMyBP-C and titin from the M-band to the C-zone. **a,** Cryo-ET map of the thick filament, spanning from the M4’ line to the C-zone. **b,** Cryo-ET map of the thick filament. with myosin tails removed for visual clarity. **c,** “Unrolled” depiction of the filament, revealing the spatial organization of the myosin heads. Position and arrangement of myosin heads from the P-zone (crowns P1, P2, and P3) to the 9^th^ stripe of cMyBP-C (from crown A1 to A28). **d**, The complete crowns’ nomenclature is indicated in. **e,** Side view of the “unrolled” filament having the center of the thick filament to the right side. **f,** “Unrolled” depiction of the filament, revealing the spatial organization of the six titin and 27 cMyBP-C molecules. **g,** Atomic models of the two titin chains and the 9 cMyBP-C within a single asymmetric unit**. h,** Domain map of titin. The pseudo-helical symmetry in the C-zone is dictated by titin’s SRs, indexing the crowns in a repeated sequence of three types (crown 1, 2, and 3). **i,** The structural arrangement reveals the 9 stripes of cMyBP-C CTD, anchored on the thick filament in proximity to the interphase of different C-type SRs.

**Fig. 3.**
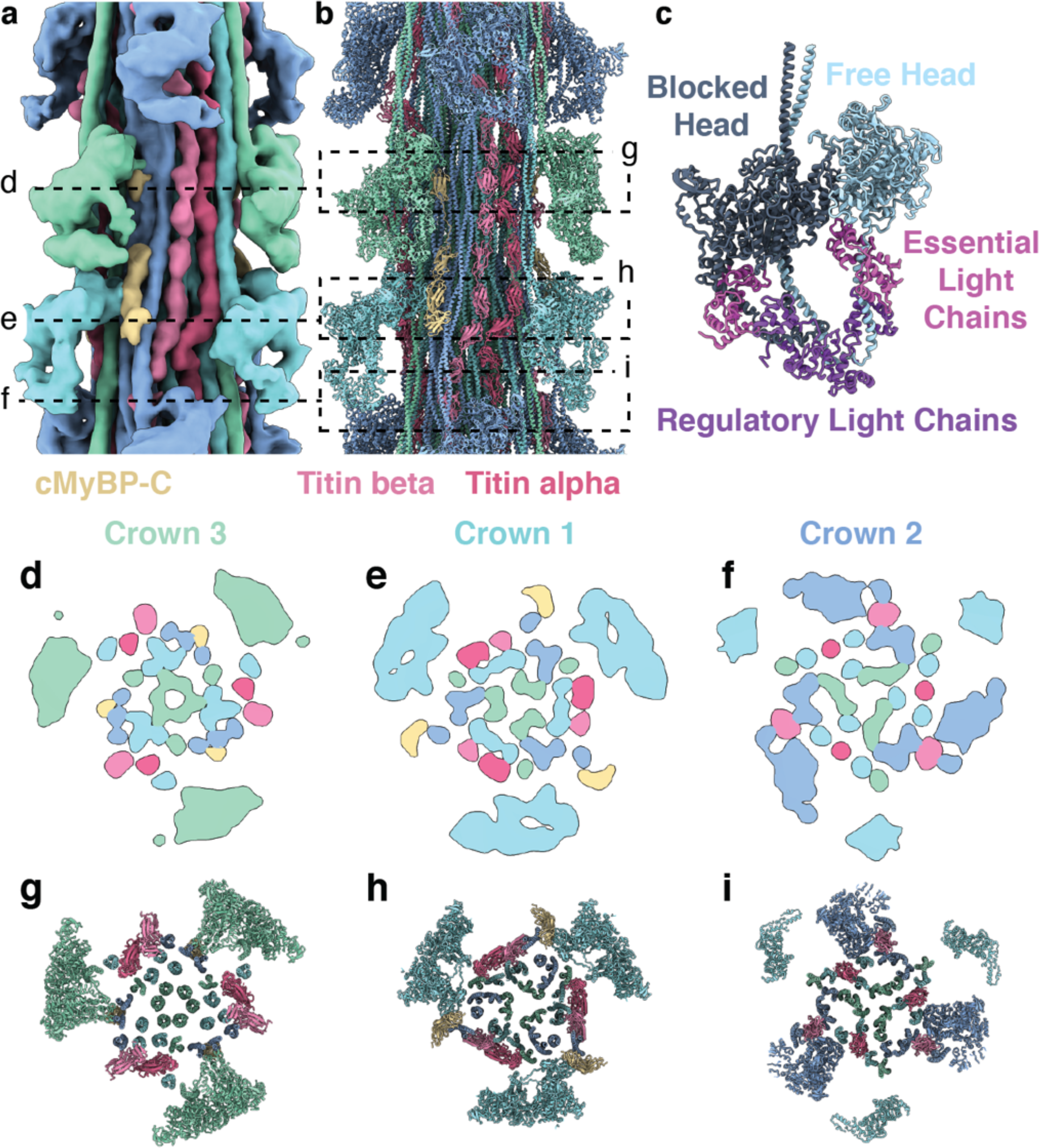
Structural model of the C-zone. **a,b,** The 3D reconstruction (**a**) and atomic model (**b**) of the C-zone, from cMyBP-C stripe #4 to stripe #9. The volume was colored according to its atomic model (**b**). **c**, Model of the myosin double heads in the IHM, with BH and FH binding to each other. **d-i,** Cross-sections of the map (**d-f**) and model (**g-i**) are shown at the level of the 3 different crowns, providing a more detailed view of the arrangement within the core of the thick filament.

The reconstruction revealed three distinct crown arrangements of myosin (defined as crowns 1, 2 and 3 from M-band to Z-disc direction) that form the outermost layer of the thick filament (Fig. 3d-i). Their double heads interact with each other within myosin dimers in the so-called interacting heads motif (IHM)^36^. In this OFF state, one of the heads is folded back (“blocked head”) so that its actin-binding site is blocked by the other head (“free head”). This configuration has been suggested to correlate with the super-relaxed state^37, 38^, where both motors have a highly inhibited ATPase activity^36^.

The myosin tails associated with the crowns form a large bundle of coiled-coils at the center of the filament (Fig. 3d-i). Interestingly, in the C-zone, besides interacting with each other, the myosin heads only bind to their own tail and are not in direct contact with the tails of other heads. Thus, there seems to be no or little direct communication possible between them.

On the bundle of tails, there are three pairs of titin molecules that run parallel to each other but where each molecule (or chain) has a unique conformation. To uniquely identify the two chains, we named them titin alpha and titin beta (Fig. 3a,b). While the heads of crown 1 and 3 do not interact with titin, the free head of crown 2 binds to the 6^th^ domain of the titin C-type super-repeat (Fig. 3,f,i), as it can also be observed at cMyBP-C stripe 1 (Extended Data Fig. 5g-i).

Importantly, the structure also demonstrates that cMyBP-C binds with its domains C7 to C10 to an array of myosin tails (Fig. 4,f-h, Extended Data Fig 6a), thus providing visual confirmation to biochemical evidence that started accumulating half a century ago^39–41^.

**Fig. 4.**
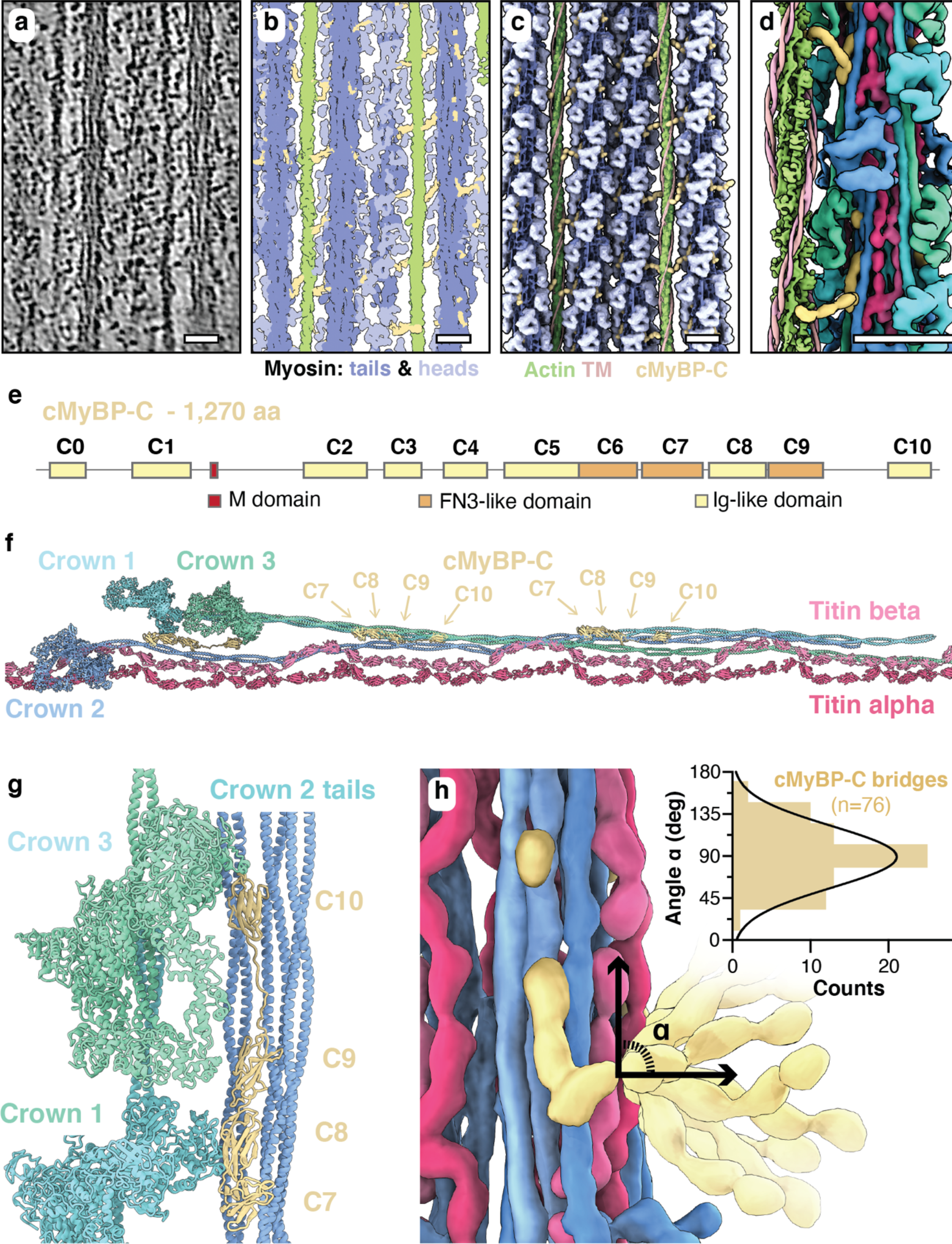
cMyBP-C forms links between the thick and thin filaments. **a**, Tomographic slice (0.94 nm thickness) of the C-zone in the relaxed state. **b**, Tomogram segmentation of a 9.4 nm slab from the same region as in (**a**), depicting the thin filament (light green), the thick filament core (blue), the myosin heads (light blue), and cMyBP-C (yellow). **c**, A 3D model of the sarcomere organization in the same slab as in (**b**), showing the flexible cMyBP-C links, spanning 3 to 4 globular domains, linking thick and thin filaments. **d**, Close-up view of the connection of two consecutive cMyBP-C to the same thin filament. **e**, Domain map of cMyBP-C. The C-terminal domains from C7 to C10 bind to the thick filament, the central domains C3 to C6 form the link, and the N-terminal domains bind to the thin filament. **f**, Structural model of the CTD of cMyBP-C, depicting how it binds to the thick filament core by multiple interactions with crown 2 tails. **g**, Schematic representation of the cMyBP-C bridging region. Flexibility is shown together with a plot of the angular distribution of the bridge measured in the tomographic volumes. Scale bars, 20 nm.

Unexpectedly, the C-terminal region of cMyBP-C also directly interacts with the IHMs. Specifically, domain C10 binds to the free head of crown 3 while domain C8 forms broad electrostatic interactions with the free head of crown 1 (Fig. 4g,h - Extended Data Fig. 5c,e). The atomic model (see Methods) indicates that this interaction likely involves loop 4 and the cardiomyopathy loop of myosin (Fig. 3b,4g, Extended Data Fig. 5c). These unforeseen interactions of the cMyBP-C with myosin motors suggest a previously undescribed direct role in the stabilization of the myosin OFF state in the C-zone and explains why cMyBP-C can stabilize the super-relaxed state^42^. This highlights a paradigm-shift for the role of cMyBP-C in sarcomere regulation and presents an interface at which unexplored pharmaceutical intervention into hypertrophic cardiomyopathy patients can be explored.

### Myosin tails exhibit distinct arrangements and pattern of interactions

Although the organization of myosin tails has been determined in insects and spiders^43–45^, to date we have no structural information to elucidate their organization in mammalian striated muscle, representing an unbridgeable gap in our understanding of the molecular mechanics of cardiac contraction. In the 43 nm repeat reconstruction of the C-zone, only parts of the tails are visible, so to visualize them in their full extent we calculated a 200 nm long helical extension starting from this reconstruction (see Methods). The resulting 3D map allowed us to elucidate the tail arrangement associated with each of the three crowns (Figure 5a). The tails of the three crowns interweave, interacting with each other and form a compact rod. The tails of crown 3 form the core of the rod and interact with each other in their C-terminal section, where the conserved 29-residue assembly competence domain is localized^46^. This domain plays a crucial role in thick filament assembly, suggesting that the myosin tails that are associated with crown 3 represent the nucleus for this process^47^.

**Fig. 5.**
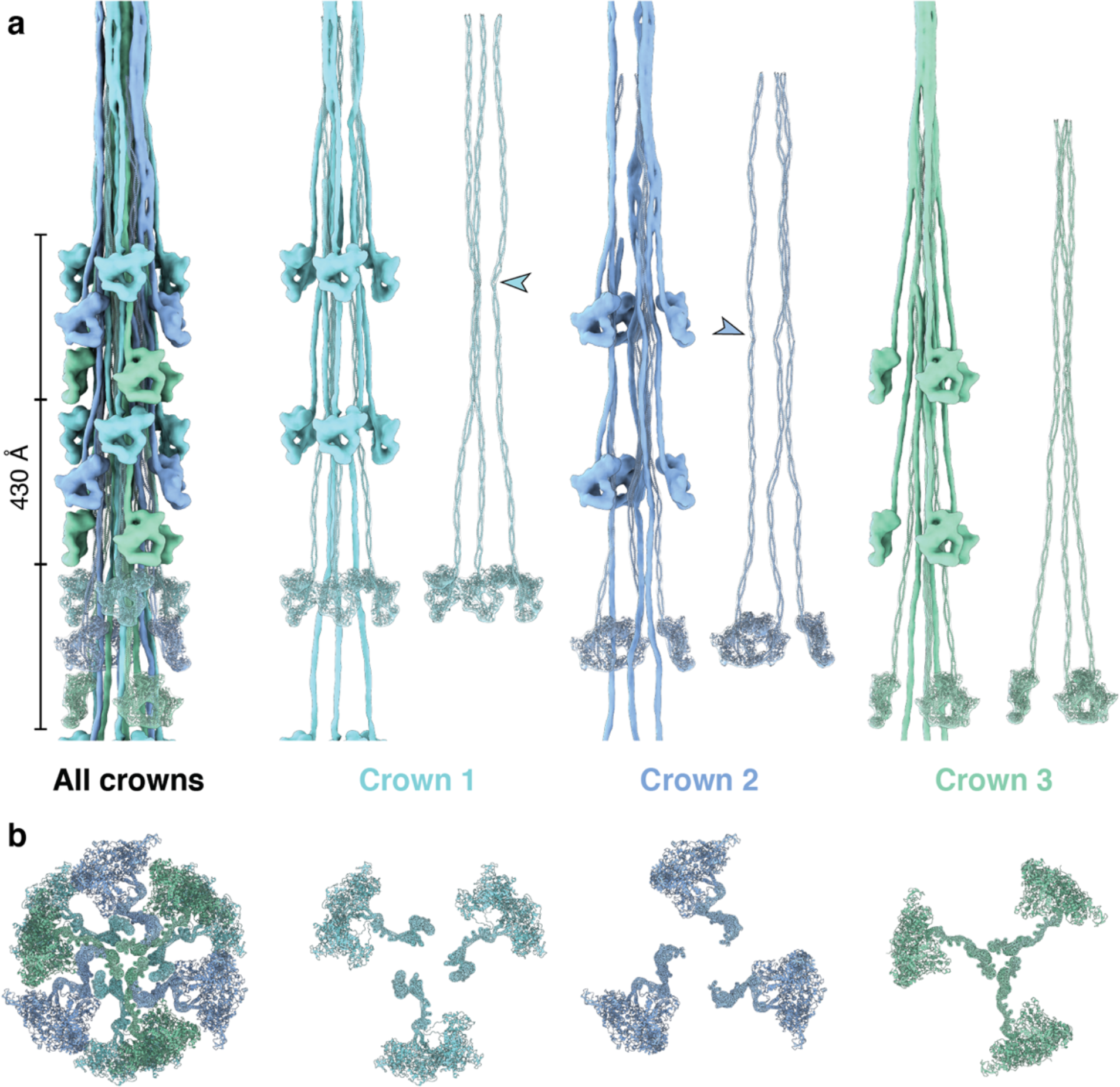
The spatial arrangement of myosin within the C-zone of the thick filament. **a-b**, Side (**a**) and top (**b**) views of a combined map and atomic models of a section of the thick filament only depicting myosins. The myosins follow a 3-fold rotational symmetry and a pseudo-helical symmetry with a 43 nm rise and 0° twist. The asymmetric unit comprises three myosin crowns, each composed of three myosin double heads. The tails form the backbone of the thick filament. The tails of crown 2 represent the outermost layer, followed by those of crown 1 and crown 3. While the tails of crown 3 run relatively straight, the crown 1 and crown 2 tails exhibit a certain degree of undulation. The arrows indicate an inflection point, common to crown 1 and crown 2 tails at phenylalanine 1449.

To quantify and compare the curved nature of the tails we calculated a sinusoidal compression percentage (see Methods) that positively correlates to increases in tail curvature and resulted in 3.05%, 4.44%, and 2.53%, for tails from crowns 1, 2 and 3, respectively. Notably, while the tails bear a unique inflection point at different locations, crown 1 and 2 share a strongly pronounced kink at the phenylalanine residue in position 1449 (Fig. 5a) that does not correspond to any skip residues predicted in the coiled-coils sequence and is not present in non-mammalian myosin II^48^, possibly revealing a peculiar characteristic of the mammalian myosin-II tail.

Within the C-zone, the myosin tails have different degrees of interaction with titin. The tails associated with crown 3 have only minimal interaction while those of crown 1 bind to either titin alpha, titin beta, or both for its entire length (Extended Data Fig. 7b,d). Tails associated with crown 2 only bind to titin beta and mostly through their S2 domain (Extended Data Fig. 7c). Taken together, these data show that each crown within the cardiac thick filament exhibits a distinct arrangement of its tails and pattern of their interactions that are likely orchestrated and tuned by the titin chains, although the reverse process would also be conceivable. This suggests the existence of specialized roles for the myosins of each crown in length-dependent activation and contraction. The curved nature of the tails of crown 1 and 2 and their direct interaction with titin supports the observation that both titin chains not only act as counters of myosin molecules (as a molecular ruler), but can sense and relay muscle load, transducing a signal that might promote transition from super-relaxed to the disordered-relaxed state of the upstream crowns towards the M-band.

### Variability of myosin crowns in the P- and C-zones

To better understand whether there are differences between myosin heads along the filament, we analyzed our combined reconstruction, where we resolved 93 myosin double heads, spanning from the P-zone layers (P1, P2, and P3) to the C-zone (from A1 to A28) (Fig. 2,a,e). We observed that all myosin heads are in the IHM state, confirming that all 31 layers are in the OFF state. The pseudo-helical arrangement of “crown 1, 2, 3” pattern, which we observed from cMyBP-C stripe 4 to 9 is preserved throughout the C-zone. However, in the regions closer to the M-band there are noticeable structural differences (Fig. 2,a,e, Extended Data Fig. 8,b-d).

To quantify these variations, we calculated the orientations (Euler angles), the azimuthal angle, axial positioning, and the radial distance for each crown (Extended Data Fig. 8,b-d). The heterogeneity in rise, twist, and angular orientation of myosin is particularly pronounced from crowns P1 to A9 while a more regular pattern emerges from A10 (Extended Data Fig. 8,b). This is exemplified by crowns A5 and A8: although they localize in canonical “crown 2” layers, these crowns have a much wider radial distance that makes them the closest to the thin filaments after P1 (Extended Data Fig. 3d). Moreover, their free heads are projected further outwards (Fig. 5a-e), as documented by their strongly negative beta angle (Extended Data Fig. 3d). This variability suggests that these crowns might have unique capacities in terms of strain susceptibility and activation rate. In particular, as they are solely stabilized by a weak interaction with titin, crown 2 appears likely to be the first responder in length-dependent activation, with exceptional layers such A5 and A8 being the most representative cases.

Taken together, although all myosins in our reconstruction are in the IHM state, their stabilization varies and relies on a variety of interactions, depending on their specific molecular context, that involves binding sites on myosin S2, titin, and cMyBP-C (see also below). The vast majority of these interactions seem reliant on the flexible loops in the free head: loop 4 and cardiomyopathy loop. Conceivably, this organization along the filament is indicative of an ability for the myosin heads within the thick filament to be fine-tuned by regulatory mechanisms of muscle contraction within the thick filament in adaptation to specific physiological conditions^49^.

### Titin organization in the P- and C-zone

Despite its essential role in sarcomere assembly and function, the detailed structural arrangement of titin has remained hitherto unknown^1, 21^. We used our combined reconstruction, spanning from the P-zone layers (P1, P2, and P3) to the C-zone (from A1 to A28) to build an atomic model of titin in which we could assign all the density to the corresponding titin domains. This allowed us for the first time to describe the molecular organization of titin from the C-zone (specifically from C-type super-repeat 2) to the M-band (Fig. 5b,e-h, Extended Data Fig. 7a) and reveal its interplay with the other thick filament components. Importantly, no structural model for titin organisation in the thick filament has been previously determined, representing a distinct lack of molecular information of fundamental importance to understanding both sarcomere function and effects causative of disease.

Surprisingly, while the three titin alpha chains run along the entire length of the thick filament and reach to the P-zone of the opposite half-sarcomere, titin beta is interrupted abruptly at myosin crown A1 (Fig. 2b,e-h). Thus, there are six chains in the C-zone, but only three in the P-zone. Consequently, only three chains enter the M-band from each side of the thick filament, resulting in only six chains of titin in the M-band instead of twelve.

In the C-type super-repeats, titin alpha and beta run alongside each other with only sparse and presumably weak interactions, resulting from the molecular contacts of titin with myosin tails. In both the P- and C-zone, titin chains show a spring-like arrangement, with recurring curved indentations following a 3-4-4 pattern that matches the alternating Fn3 and Ig-like sequences found in titin’s super-repeats^50, 51^ (Extended Data Fig. 6a-d). Beside this common motif, we observed a certain degree of variability between different C-type super-repeats (Fig. 2b,e-h). These variations are mostly accounted for by the disorganized linkers scattered between domains all along the structure and are more pronounced on titin beta. The super-repeat domains from 4 to 8 are those showing the highest conformational variability in titin beta. These domains interact with the S2 domain of crown 2 (Extended Data Fig. 5b,g-j) and their variability is reflected in the heterogeneous configuration of crown 2 IHMs (Extended Data Fig. 8b-d) as already discussed.

Missense mutations in the A-band portion of titin which includes the C-zone have been linked to multiple, often severe myopathies and cardiomyopathies^52, 53^. A hotspot for variants linked to hereditary myopathy with early respiratory failure (HMERF) is A150/Fn3-119^52^. The proximal S2 domain of crown A5 sits on this domain, suggesting that a functional impairment of this unique crown might result in this specific muscle disease (Figure 2, Extended Data Fig. 5g-j).

In the P-zone, which is the transition zone between the A- and M-band, the pattern of the titin C-type super-repeat is replaced by 7 domains (from A164 to A170) that lead to the titin kinase (TK) domain (Fig. 2b,e-h). Following the titin alpha chain in our reconstruction, we identified a bulkier region behind the P1 crown that fits the size, the shape, but not the expected position of the TK (Extended Data Fig. 9b). As this position disagrees with that previously assigned at ∼52 nm from the M1-line by immuno-electron microscopy using a monoclonal IgM anti-TK antibody^54^, we generated a new affinity-purified TK antibody and used it to ascertain the TK position by super-resolution microscopy in mouse cardiac and mouse and rabbit psoas myofibrils (Extended Data Fig. 9g). The position of the TK domain thus determined was 78 nm ± 8 nm from the M1-line, consistent with the titin density at the P1 crown. TK interacts with myosin’s S2 domain, and is closely connected via its C-terminal regulatory tail to the adjacent m1 domain, as has been previously proposed^55, 56^. Interestingly, the titin m1 domains interact with the S2 domains of the myosins in crown P2, while the subsequent titin m2 and m3 domains are wrapped around the myosin tails associated with the P1 crown (Extended Data Fig. 9b).

In the C-zone, the titin beta chain follows the same register of titin alpha, however, after domain A154, in proximity of crown A2, its organization changes to a bowtie-shaped structure and its chain is absent after crown A1 (Extended Data Fig. 9d). On top of the A1 crown, we identified another bulky density reminiscent of the TK-m1 configuration. However, the limited resolution does not allow a clear identification of domains. We propose that the bowtie-shaped structure might correspond to 13 titin domains (from A158 to A170), connecting the rest of titin beta to the beta TK-m1 domain (Extended Data Fig. 9d) and that titin beta terminates its run along the thick filament on the A1 motor domains, perhaps after proteolytic cleavage by calpain 3 (CAPN3) downstream of titin m1^57^. Thus, the TK domains of titin alpha and possibly of titin beta would localize at 77 and 105 nm from the centre of the M-band (in contact with crowns P1 and A1, respectively) interacting with their S1 and S2 domains in privileged positions that could couple myosin head activation to mechano-sensing by TK. Our super-resolution microscopy analysis does not detect an additional signal at crown A1, this might result from a less accessible position of the epitopes to the antibodies. Alternatively, the bowtie-shaped structure could also represent a so-far uncharacterized protein complex interacting with titin beta.

### The molecular organization of the cardiac M-band

The M-band, comprising 6 transverse “lines” (M1-M6), is a complex network of structural proteins, metabolic enzymes and proteostatic machinery components^1, 4^. Proteins in this network are reported to anchor the M-region to the sarcolemma, stabilize the myosin filament hexagonal lattice, and to serve as a signaling scaffold that integrates mechanical forces, energy balance and protein turnover^1, 4^. Our reconstruction of the thick-filament M-region shows a twofold mirror axis centered around the M1 line, perpendicular to a threefold rotational axis. We could clearly identify the myosin crowns in the P-zone and also follow the tails from the innermost 4 crowns, that run antiparallel to each other, thus confirming previous indications^5^ that the P1 tails terminate in proximity of the P1 crown from the opposite polarity (P1’) (Fig. 6a,c).

**Fig. 6.**
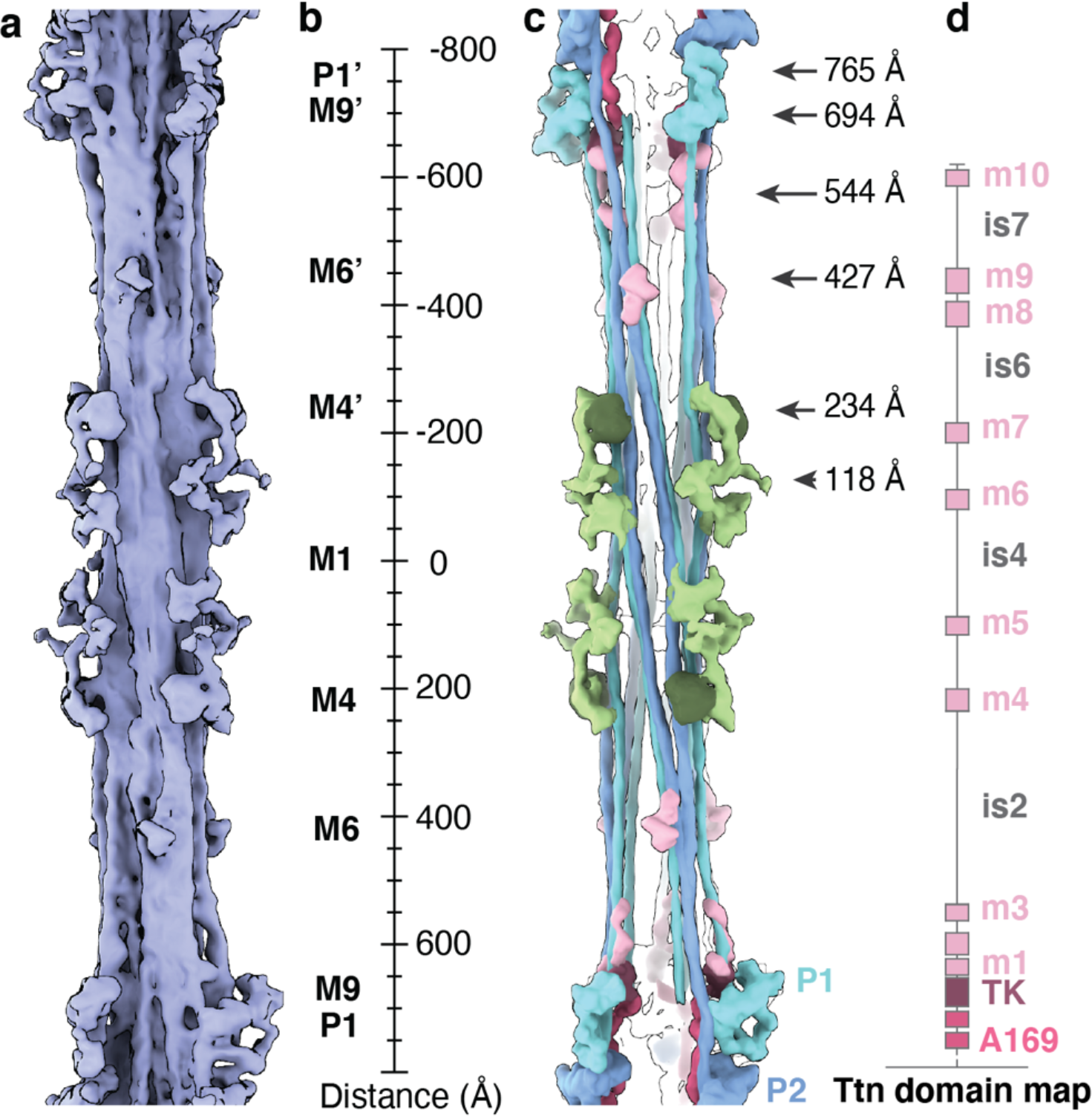
The layout of the M-band. **a**, Density of the thick filament in the M-band region. The overall structure shows a twofold rotational axis perpendicular to a threefold rotational axis (D3 symmetry). **b,** Ruler, providing markers for the expected high-density M-lines in the cardiac sarcomere (M1, M4, M6, M9). M1 is absent in our reconstruction, likely due to the lattice organization being averaged out during the refinement process. **c,** Most clearly identifiable densities protruding from the core of the myosin tails. **d**, Domain map of the titin C-terminal region, which consists of the titin kinase domain (TK), the ten m-domains (m1-10), and seven interspacing regions (is1-7).

In addition to myosin, we observed several densities protruding from the core of the tails, reflecting M-band-specific proteins such as myomesin 1 and 2^4^, obscurin-like-1 (OBSL1^58^) titin C-terminal domains^59^ and accessory proteins like muscle-type creatine kinase (M-CK). We identified a density spanning from M-lines M1 to M4 that binds to the thick filament core at 85 Å and 230 Å from M1 and projects outward with a threefold symmetry into the hexagonal lattice. Taking into consideration its localization, arrangement and composition from ∼4 nm-long domains, we suggest this density to represent a complex of myomesin-1, OBSL1 and possibly myomesin-2 (Fig. 6a-c). Surprisingly, in our super-relaxed cardiac M-band reconstruction, we observe densities at 118 Å from the M1, protruding towards neighbor thick filaments in the lattice (Fig. 6c). However, the expected cross-connecting M-bridges, as they would be inferred from the dimeric myomesin-1 structure^60^, are averaged out during the refinement due to their flexibility.

We could not determine the location of obscurin. This is likely due to a combination of structural flexibility and the localization of obscurin at the myofibril periphery, as opposed to the central sarcomeric localization of OBSL1^58^. As described above, only titin alpha reaches the M-band region with its C-terminus. It consists of the TK domain, ten m-domains (m1-10), and seven interspacing regions (is1-7)^54^. Within each asymmetrical unit of the bare zone, at 427 Å from the high-density M-line M1, we observed a density that fits the size and shape of two Ig-like domains, possibly the adjacent m8 and m9 titin domains. In addition, there are other small densities that might correspond to m1, m2, m3 and m10. The interspacing regions are very thin and likely contain few secondary structure elements, and therefore cannot be seen in our reconstruction. In line with previous suggestions^54^, these observations support that the titin alpha M-region spans across the M-band. The identification of M-line densities in their near-native state provides a blueprint for titin m-domain localization and structural rationalization of reported biochemical interactions^4, 58, 59, 61–63^. Detailed assignment of the densities will require a combination of site-directed labeling of the domains and cryo-ET in the future.

Nonetheless, the structure of the native cardiac M-band presented here *in situ* represents an initial model of the M-band at the molecular scale. The crosslinks formed by M-band proteins are dictated by the threefold rotational symmetry of the thick filament. However, the interactions of titin with OBSL1, and OBSL1 with myomesin-1 are all in a 1:1 complex^62, 63^, posing a symmetry mismatch between these constituents and the previously expected six titins from each half-sarcomere entering the M-band. Our analysis resolved this paradox, showing that the full M-band region of titin is only present in three copies in each half-sarcomere, directing the assembly of the six observable M-links (three in each half-sarcomere).

### cMyBP-C forms links between thick and thin filament

In our reconstruction of the thick filament, we identified that the C7-C10 domains of cMyBP-C bind to an array of three consecutive tails associated with crown 2 in 9 regions of the C-zone (Fig. 2a-b,f-g,i)^24, 64^. C7-C10 run longitudinally along the filament: for each cMyBP-C, C10 is in the M-band direction and C7 is oriented toward the Z-disc, spanning a 16 nm region at the interphase between titin C-type super-repeats (Fig. 2f-i). Initial biochemical data suggested that the confinement of cMyBP-C to the C-zone might depend on a direct interaction between the C7-C10 of cMyBP-C and the titin C-type super-repeats^13^. We do not observe any direct interactions between the two proteins in the relaxed state. Instead, we clarified that the C-terminal domains bind to a specific configuration of myosin tails that emerges in the C-zone as a complex interplay between myosin tails and titin C-type super-repeats (Fig. 4f,g, Extended Data Fig. 5b-d). Specifically, C7-C10 bind to regions that emerge from an array of tails associated with crown 2 belonging to three consecutive C-zones (Fig. 4f,g) via strong electrostatic interactions (Extended Data Fig. 5d,f). This configuration also resolves the mysterious absence of cMyBP-C in the titin region of titin super-repeat 1, as the site where cMyBP-C would bind is in this case composed of tails of the D-type super-repeat 6, which we have not resolved in our structure, but which have likely a different conformation.

In the cMyBP-C C-terminal region, C7 is the last clearly resolved domain and appears to act as a pivot point for the remaining domains that reach away from the thick filament (Fig. 3a). The bridging region of the protein indeed showed great flexibility and only had weak densities (Fig. 4h - Extended Data Fig. 10a,b). Although stripes 1, 2 showed biases towards a specific lattice organization the cross-sections analysis at the different stripes revealed that the MyBP-C N-terminal region can bind to either of the two neighbouring thin filaments (Extended Data Fig. 10a,b,c). Remarkably, cMyBP-C stripe 2 shows some unique features (Extended Data Fig. 11): the free head of crown A7 is not stabilized by an interaction with cMyBP-C C8, as it is observed in all the other stripes, instead its IHM is stabilized by interaction with myosin light chains of the neighbor crowns.

To validate this pliability of cMyBP-C links, we back-plotted the structures into the reconstructions of the tomographic volume and manually traced the flexible globular domains connecting thick and thin filaments (see Methods) (Fig. 4a-d, Supplementary Video 3). We observed the presence of several cMyBP-C links in the C-zone, suggesting that the N-terminal region can indeed interact with the thin filament. Despite our efforts, sub-tomogram averaging did not resolve any clear arrangement of the N-terminal region on the thin filament, indicating a high heterogeneity in these interactions, in agreement with previous reconstructions from isolated components^65^. Our segmentation exposed the presence of a tether formed by 3 to 4 domains that blend in the signal of the thin filament with high degrees of flexibility (Fig. 3h). Of note, the thin filament with bound N-terminal cMyBP-C domains appears in the OFF-state, suggesting that cMyBP-C, at physiological phosphorylation levels, and in the absence of myosin cross-bridges, is not a thin filament activator.

cMyBP-C represents a crucial site of cardiomyopathies and has thus far remained an enigmatic component of the sarcomere, with its function in modulating myosin activation being both proposed but not well understood. Collectively, our analysis of cMyBP-C suggests multiple roles in sarcomeric mechano-signaling. On the one hand, it is reasonable to speculate that under thick filament strain, the sinusoidal path of crown 2 tails can experience a minimal but significant stretch, resulting in the alteration of the binding sites for the cMyBP-C C-terminal region and perturbing its interactions with the free heads. As such, the interactions of the C-terminal domains with the IHMs of crown 3 and 1 would represent a transducer for tension and stretch, independently of thin filament positioning. On the other hand, the crosslinks to the thin filament can sense the stretch on and sliding of the thin filament and relay the tension to both the C-terminal and N-terminal domains of cMyBP-C, while domain C1 and C0 might extend further to bind the myosin light chains on the thick filament surface, either in cis or trans.

### Conclusions

Our study provides important insights concerning the fundamental organization and regulation of cardiac thick filaments in vertebrates. Our cryo-ET structure of the sarcomere in the relaxed state together with our previously determined structures of the sarcomere in the rigor state^20, 31^, provide fundamental insights into muscle architecture, its molecular complexity, intricacy of regulation and sophistication. The structure of the native thick filament provides a three-dimensional representation of how the myosin heads and tails, myosin-binding protein C, and titin are organized in different areas of the filament, explaining how these components can interact during muscle contraction. It also reveals distinct differences in how certain domains of the sarcomere are organized when compared to tradition reconstituted approaches and classical EM methods.

Our analysis revealed a plethora of conformations and stabilization mechanisms for the IHM, which are region-dependent and rely on various interactions with titin m1 domain, cMyBP-C C-terminal region, titin SRs, other myosin tails, and myosin light chains. These findings support a model of a thick filament that is capable of sensing, integrating, and modulating responses to numerous signals such as beta-adrenergic-dependent phosphorylation, Ca^2+^ concentration, length, and mechanical load. In addition, we revealed the organization of titin along the thick filament and the genuine position of its TK domain not in the M-band but juxtaposed at the interface between the M-band and A-band. Such a localization indicates that the role of TK in regulating the sarcomere may be more tightly coupled to the state of the myosin heads than previously thought. The six titin chains probably influence the arrangement and pattern of interactions of each myosin tails. Thus, it is understandable how titin acts as a scaffold and a molecular ruler for myosin assembly during sarcomerogenesis, as has been previously suggested^66^. This is an interesting reflection of the protein nebulin found in the thin filament^31^. Furthermore, the close interaction and spring-like appearance of both titin and the myosin tails and their connection to myosin heads make a compelling case for a load-dependent activation mechanism of the thick filament promoting the transition from the super-relaxed to the disordered relaxed state.

We were able to observe biologically significant structural variability along the thick filament, which is a crucial aspect needed for understanding sarcomere function and regulation. Static structures in isolation tell us little about the interplay and dynamics of multiple, highly regulated myosin heads interlinked to one another. These important details would have been lost with most structural biology methods and demonstrate the merits of cryo-ET in visualizing this variability in the native state. The MyBP-C links connecting thin and thick filaments are another example of the unique power of cryo-ET. Finally, the insights gleaned through our thick filament structures lay a framework for understanding muscle development, function and muscle disorders.

## Methods

### Myofibrils preparation and vitrification

Demembranised left-ventricular mouse myofibrils were prepared as previously described^31^. The myofibrils were collected by centrifugation at 3000 g for 2 minutes at 4°C, followed by two washes with pre-relaxing buffer (100 mM TES pH 7.1, 70 mM KCl, 10 mM reduced glutathione, 7 mM MgCl^2^, 25 mM EGTA, 20 μM mavacamten, 5% dextran T500). To prepare for plunging, the pre-relaxing buffer was replaced with final relaxing buffer (pre-relaxing buffer plus 5.5 mM ATP). The relaxed myofibrils were then frozen onto Quantifoil Au R2/2 SiO^2^ 200 mesh grids using a Vitrobot (Thermo Fisher). The myofibril suspension was incubated on grid at 25°C and 100% humidity for 30 seconds, blotted for 30 seconds from the opposite side of the carbon layer, and plunged into a liquid ethane-propane mixture.

### Cryo-FIB milling and electron cryo-tomography (cryo-ET)

The preparation of lamellae for cryo-ET data acquisition was performed by cryo-FIB milling using an Aquilos 2 cryo-FIB/SEM with a cryo-shield, according to previously described protocols ^20, 30, 31^ aiming for lamellae with a final thickness of 180 nm (range from 90 to 250 nm). The data acquisition was carried out with a Titan Krios transmission electron microscope (Thermo Fisher), fitted with a K3 camera and an energy filter (Gatan). The acquisition of overview images of myofibrils in the lamellae was done at a nominal magnification of 6,700× to identify the C-zones regions. The average sarcomere length was 2.33 μm (S.D. = 0.16 μm, N = 29).

The tilt series were acquired targeting the C-zones at 81,000× nominal magnification. The pixel size was calibrated to 1.146 Å using the 143.3 Å peak in the fast Fourier transform of the final thick filament reconstruction (from M-band to C-zone) (Extended Data Fig. 4a,c). A dose-symmetric tilting scheme^67^ was applied during acquisition with a tilt range of −50° to 50° relative to the lamella plane at 2.5° increments. The sample was subjected to a total dose of 120 to 160 e^−^/Å^2^. Tilt series were acquired between −3 and −6 µm defocus. All images and a total of 89 tomograms were acquired using SerialEM^68^.

### Tomogram reconstruction and particles picking

Motion correction and contrast transfer contrast (CTF) estimation was performed in Warp^69^, tilt series alignment was performed in IMOD^70^. Final tomogram reconstruction and sub-tomograms extraction was performed in Warp. After binning the tomogram to a pixel size of 0.92 nm and low-pass filtering them at 60 Å, we used crYOLO^71^ to pick and trace both the thick and the thin filaments (Extended Data Fig.1).

### Thin filament processing pipeline, model building, and visualization

The traced thin filaments were re-sampled with an inter-segments distance of 18 Å, leading to the extraction of 365,971 sub-tomograms with a box size of 293.5 Å (128 pixels, binning 2). Each sub-tomogram was rotated to orientate the thin filaments parallel to the XY plane using the prior angles from the tracing. Then its central slab of 100 slices was projected and used as an input for two-dimensional classification^72–74^. The classes that did not show a clear presence of thin filaments were discarded and the remaining segments were processed in RELION 3.1^75, 76^. The initial helical reconstruction led to 14.3 Å resolution map (0.143 FSC criterion) with 27.4 Å rise and −167.2° twist (Extended Data Fig.1). After removing the duplicated particles using a customized script, 100,447 sub-tomograms were further refined with two different masks that either covered the entire density of the thin filament, or only included the F-actin density. The full thin filament (F-actin and tropomyosin) was refined with helical reconstruction and reached a resolution of 8.2 Å while the refinement of F-actin alone resulted in an 8.3 Å resolution map. The two maps were aligned in ChimeraX^77^ and the individual chains from the PDB model 6KN7^78^ were placed in the density with rigid body fitting. The model of tropomyosin’s coiled-coils was improved with Namdinator, using automatic molecular dynamic flexible fitting (MDFF)^79^. The final composite map (Extended Data Fig.3a) was created by combining F-actin from the F-actin alone reconstruction and tropomyosin from the full thin filament reconstruction using “color zone” and “splitbyzone” functions in ChimeraX.

### Thick filament processing pipeline

The traced thick filaments were re-sampled with an inter-segments distance of 130 Å, leading to the extraction of 67,492 sub-tomograms with a box size of 1280 Å (160 pixels, pixel size 8 Å). 2D classification was performed similarly to the thin filament processing, using a central slab of 400 Å, resulting in 37,118 high-quality sub-tomograms. 3D classification with refinement and helical reconstruction (430 Å, 0° twist) resolved 4 classes that showed a different orientation of crown 2. Class A showed a “projected” IHM, class B had a mixture of conformations resulting in fuzzy density for crown 2 IHM, and class C showed a “retracted” IHM (Extended Data Fig.1b). The refined coordinates were used to individually re-extract the 3 classes from Warp, using a box-size of 144 pixels and a pixel size of 4 Å. The individual classes were refined with RELION and their coordinates were mapped back into the tomograms using ArtiaX^80^. Class A, which was later resolved as the segment from crown A8 to A12 (cMyBP-C stripe #2) (Extended Data Fig.1g), showed a unique distribution within the sarcomere. This was used as the initial anchor point to obtain the 3D coordinates of the particles located at an axial distance of −43 nm and +43 nm (Extended Data Fig.1f,h). Through an iterative process of 3D refinement, multi-reference 3D classification, particles back-projection, and axial shift calculations, we resolved 8 structures of the thick filament, spanning from M-band to C-zone (Extended Data Fig.1). The resulting 3D maps showed high variability in resolution within different regions of the same map leading to over-sharpening of the more flexible regions, we therefore resorted to LocSpiral^81^ to improve map interpretability. All reconstructions showed a three-fold rotational axis and were therefore refined with C3 symmetry. The M–band reconstruction further revealed two orthogonal two-fold rotational symmetry axes that intersect at the three-fold axis at an angle of 60° and was later refined with D3 symmetry.

To build the final composite map, the power spectra of the reconstructions were normalized with *relion_image_handler*^76^ and a soft cylindrical mask of 15 px (∼ 60 Å) was applied to all maps. The filtered reconstructions were aligned in ChimeraX (*fit in map*) and the final map was created by merging the densities of the different segment, using the maximum value at each voxel (*volume add*, *volume maximum* in ChimeraX). To obtain a continuous and homogeneous density that we could use to trace the myosin tails, we took our 18 Å reconstruction of the last 5 stripes of cMyBP-C and extrapolated a 200 nm long helix with *relion_image_handler* (430 Å, 0° twist)^76^.

### Thick filament’s model building and visualization

The model of the thick filament was built using a combination of previously available models and AlphaFold2 predictions^35^. For the model of myosin-II’s, we used the IHM of human beta cardiac heavy meromyosin (PDB entry 5TBY^34^) while the tails were predicted in AlphaFold2 using the amino acid sequence of *Mus musculus*‘s MYH7 (5 segments of ∼250 amino acids with ∼20 amino acids overlap). Similarly, the C-terminal domains of cMyBP-C were predicted in AlphaFold2 using the last 590 amino acids of *Mus musculus*‘s MYBPC3. The region of titin from domain A101 to m3 (amino acids 24,760 – 32,350) was submitted for prediction as multiple entries (each ∼950 amino acid) with overlapping terminal domains. The models were initially built in the map using rigid body fitting and their final organization was later adjusted with multiple rounds of MDFF in Namdinator^79^, starting with 40 Å low-pass filtered densities and gradually using the higher-resolution maps. The models spanning from crown A18 to A28 were obtained by cloning the model spanning from crown A15 to A17. For the visualization in Fig. 2a, we resort to ChimeraX functions “*colorbyzone*”, “*splitbyzone*”, and “*Gaussian filter*” with standard deviation = 3. The depictions in Fig. 2c,e,f were obtained using Chimera “*unroll*” function on the structure of all components individually.

### Position and orientation of myosin’s crowns

To describe the differences in the IHMs across our entire reconstruction, for each crown we calculated the cylindrical coordinate (azimuthal angle, radius, and z-axis height) and extracted radial distance, rise, and twist (Extended Data Figure 8d). Additionally, we calculated the Euler angles (alpha, beta, and gamma angles) to better describe their 3D arrangement (Extended Data Figure 8c). To do this, we treated each IHM as a triangle lying on the surface of a cylinder (the thick filament). The coordinates of the three vertices were obtained from the atomic coordinates for the alpha-carbons of three residues: chain A residue 180, chain B residue 180, and chain A residue 830 (Extended Data Figure 8a). Cylindrical coordinates were obtained from the triangle’s centroid, while the Euler angels are relative to a reference triangle that lies on the lateral surface of the cylinder and has its top side parallel to the cylinder’s plane and tangential to its radius.

### Sinusoidal compression percentage

To quantify the curvature of the myosin tails, for each tail, we first obtained the atomic coordinates of the alpha-carbons for the two amino acids chains in the coiled-coils. We then traced a new three-dimensional curve running through the central points between each alpha-carbon couple. For each curve segment, we calculated the sinuosity S by the ratio of length of curve C to the Euclidean distance between the ends L:

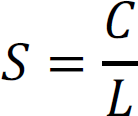

The sinusoidal compression percentage (SCP) is then given by:

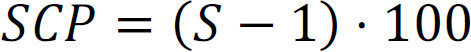

### Tomogram segmentation and cMyBP-C links

To describe the 3D organization of the sarcomere components we selected two representative tomograms (Fig. 1a, 4a, Supplementary Videos 1 and 2) and denoised them using cryo-CARE^82^. With a customized script, we mapped back each sub-tomogram using a binary mask of their corresponding structure, matching the coordinates and orientations obtained from the 3D refinement. The resulting binary MRC files were imported in Dragonfly^83^ and used as a template for pseudo-segmentation of the tomograms. The resulting label-layers were manually validated by inspecting each tomographic slice and further tracing the flexible components that were averaged out during the refinement (i.e. the cMyBP-C links from thick to thin filament).

After clearly identifying and segmenting 76 cMyBP-C links in our tomograms, we measured the angle that the link formed relative to the thick filament z-axis, using the position of the C7 domain as pivot point. The angular distribution was plotted in GraphPad Prism.

### Antigen expression and purification

A fragment of TK encompassing human TTN transcript variant-IC (NM_001267550.1), residues 33812 – 34076, was expressed in *E. coli* BL21 [DE3] cells in fusion with an N-terminal His^6^ tag. The insoluble fragment was extracted from inclusion bodies with 8 M urea, 50 mM potassium phosphate pH 8.0, 0.5% Tween 20 (buffer B) by sonication with a Branson sonifier microtip on ice. Insoluble material was pelleted at 15 krpm, 20 min in a SA 600 rotor (Sorvall) and the soluble supernatant applied to a Ni-NTA column equilibrated in buffer B. After washing the column as above in buffer B, bound protein was eluted with 250 mM imidazole in buffer B and equilibrated stepwise against 6, 4, 2 and 0 M urea in 40 mM HEPES buffer pH 7, 50 mM NaCl, 4 mM DTT and 0.1 % Tween 20 (buffer C). Insoluble precipitate was spun down and soluble protein further purified by gel filtration purification on a Pharmacia Superose 12 column equilibrated in buffer C. The purified kinase fragment was used for commercial rabbit immunisation, and serum was collected after 3 booster injections.

### Cloning, expression and purification of rat titin A170-Kinase

For affinity purification, a soluble TK construct, A170-kinase, was used. The sequence encompassing the A170 (Fn3) and kinase domains of rat titin (XM_008775521.1 residues 31897 – 32344) was cloned into a modified pCDFDuet vector containing an N-terminal His-tag, expressed in *E. coli* strain BL21 [DE3] using standard protocols and purified by nickel affinity and size-exclusion chromatography according to Rees, et, al.^53^.

### Antigen coupling and affinity purification of anti-titin kinase antibody

1 mg purified A170-Kinase was dialysed into coupling buffer (100 mM sodium phosphate pH 8, 250mM NaCl, 1mM DTT), then coupled to 2 ml NHS-activated Sepharose 4 Fast Flow slurry following the manufacturer’s instructions (Cytiva Life Sciences). Antibody affinity purification was performed using standard procedures described by Harlow and Lane^84^. Following equilibration with 10 ml PBS/0.05% Tween-20, 5 ml of the rabbit anti-kinase serum was applied to the A170-kinase – Sepharose column, which was then washed with 20 ml PBS/0.05% Tween-20, 4 ml PBS and finally 4 ml 50 mM NaPO^4^ pH 7.4, 500 mM NaCl to remove non-specifically bound proteins. Bound antibodies were then eluted with fractions of 0.5 ml 0.1 M glycine/HCl pH 3 into 1 ml 1 M Tris/HCl pH 9, with those containing protein pooled, dialysed into PBS containing 5 mM NaN^3^, concentrated to ∼0.24 mg/ml, flash-frozen in 50 µl aliquots and stored at −80°C. Specific reactivity of the purified immunoglobulins was confirmed by Western blotting against various titin fragments containing the kinase as well as control fragments.

### Super-resolution microscopy

Immunofluorescence labelling was performed on mouse and rabbit psoas myofibrils as previously described^19^, using the affinity-purified TK antibody at 1 µg/ml and Atto647N-labelled anti-rabbit IgG secondary antibody for visualization. STED was carried out on a STEDYCON (Abberior) attached to a Leica TCS SP5 ll confocal microscope. Images were recorded at a pixel size of 15 nm.

## Acknowledgements

We gratefully thank D. Prumbaum and O. Hofnagel for the assistance with cryo-EM data collection, A. Oelschläger for scripting the calculation of myosin heads orientation and R.S. Goody, F. Schnorrer, and W. Oosterheert for critical proofreading of the manuscript. This work was supported by funds from the Max Planck Society (to S.R.), the Wellcome Trust (Collaborative Award in Sciences 201543/Z/16/Z to S.R. and M.Ga.), the European Research Council under the European Union’s Horizon 2020 Programme (ERC-2019-SyG, grant no. 856118 to S.R. and M.Ga.), and the Medical Research Council (MR/R003106/1 to M.Ga. and A.L.K.). D.T. was supported by an IMPRS fellowship. M.Gr. was supported by an EMBO Long-Term Fellowship. M.Ga. holds the BHF Chair of Molecular Cardiology.

## Author contributions

S.R. designed and supervised the study. D.T. performed cryo-FIB milling, cryo-ET experiments, performed sub-tomogram averaging, analyzed data, and prepared figures. Z.W. guided the initial reconstruction of the thin filament. S.T., T.W. and M.S. wrote scripts for particle re-extraction and re-projection. Z.W., S.T. and M.Gr. optimized initial FIB-milling and cryo-ET acquisition. A.L.K. and M.Ga. developed methods and prepared myofibril samples. M.R. cloned, expressed, purified, and verified the anti-TK antibodies. P.B. performed super-resolution microscopy experiments. D.T. and S.R. wrote the manuscript with contributions from all authors.

## Additional Information

Correspondence and requests for materials should be addressed to S.R.

## Data availability

Cryo-ET structures have been deposited to the Electron Microscopy Data Bank (EMDB) under accession numbers (dataset in brackets): EMD-16986 (thin filament with masked out tropomyosin) [https://www.ebi.ac.uk/emdb/EMD-16986], EMD-16987 (thin filament including tropomyosin) [https://www.ebi.ac.uk/emdb/EMD-16987], EMD-16990 (Crowns P2-A1 from the relaxed thick filament) [https://www.ebi.ac.uk/emdb/EMD-16990], EMD-16991 (M-band from the relaxed thick filament) [https://www.ebi.ac.uk/emdb/EMD-16991], EMD-16992 (Crowns A15-A29 from the relaxed thick filament) [https://www.ebi.ac.uk/emdb/EMD-16992], EMD-16993 (Crown P1 from the relaxed thick filament) [https://www.ebi.ac.uk/emdb/EMD-16993], EMD-16994 (Crowns A11-A15 from the relaxed thick filament) [https://www.ebi.ac.uk/emdb/EMD-16994], EMD-16995 (Crowns A8-A12 from the relaxed thick filament) [https://www.ebi.ac.uk/emdb/EMD-16995], EMD-16996 (Crowns A5-A7 from the relaxed thick filament) [https://www.ebi.ac.uk/emdb/EMD-16996], EMD-16997 (Crowns A1-A5 from the relaxed thick filament) [https://www.ebi.ac.uk/emdb/EMD-16997].

Representative tomograms have been deposited under accession numbers EMD-16989 (Tomogram of sarcomere M-band to C-zone from mouse cardiac muscle) [https://www.ebi.ac.uk/emdb/EMD-16989] and EMD-16988 (Tomogram of sarcomere C-zone from mouse cardiac muscle) [https://www.ebi.ac.uk/emdb/EMD-16988].

