## Supplementary Figures for "In situ structures from relaxed cardiac myofibrils reveal the organization of the muscle thick filament"

### Extended Data

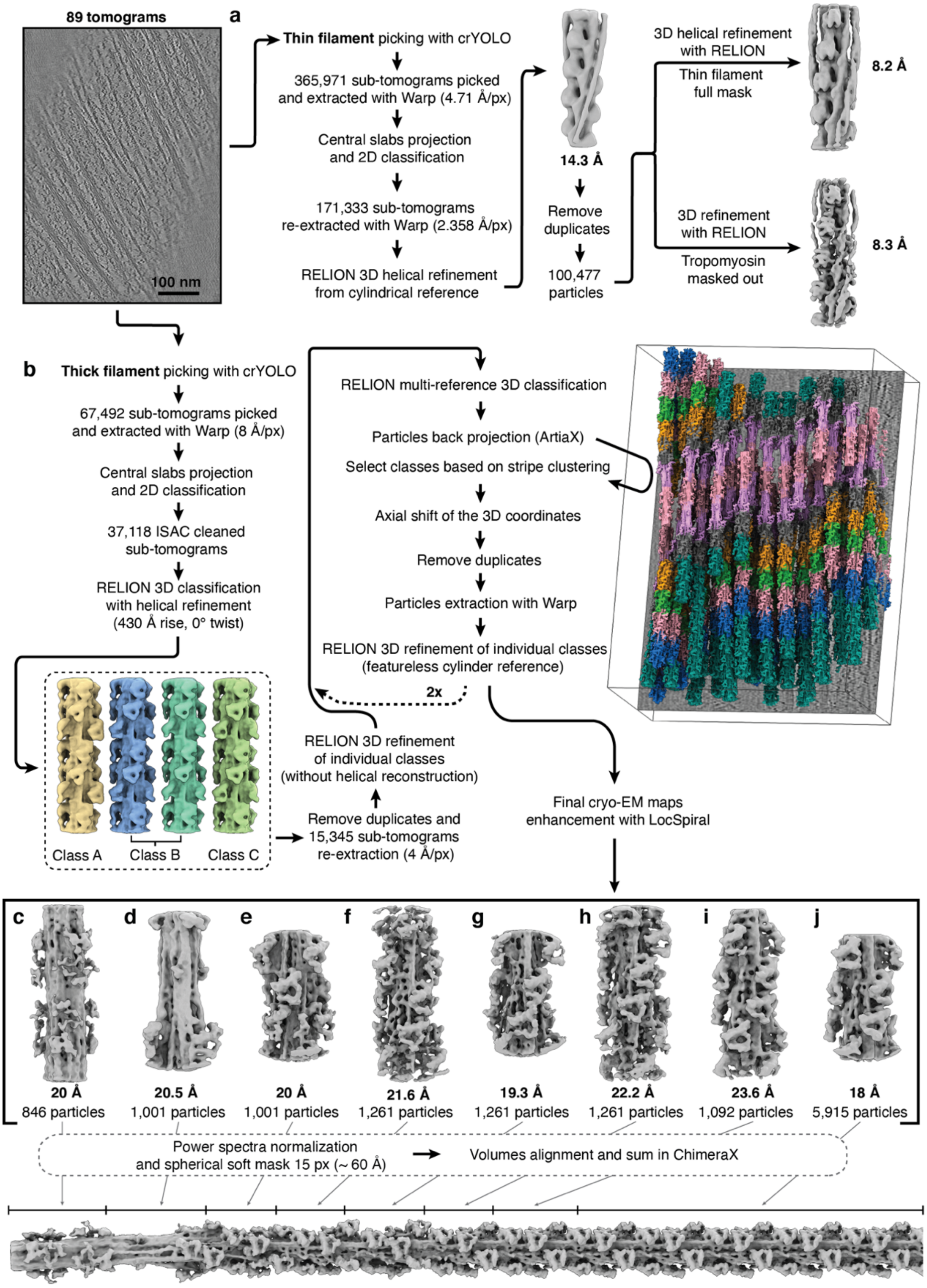


**Extended Data Figure 1** Data acquisition and processing pipeline of the thin (**a**) and thick filament (**b**). Cryo-EM enhanced maps for: M-band (**c**), A-M transition to crown P1 (**d**), from crown P2 to A1 (**e**), from crown A1 to A5 (**f**), from crown A5 to A7 (**g**), from crown A8 to A12 (**h**), from crown A11 to A15 (**i**), and the combined segments from crown A15 to A29 (**j**)

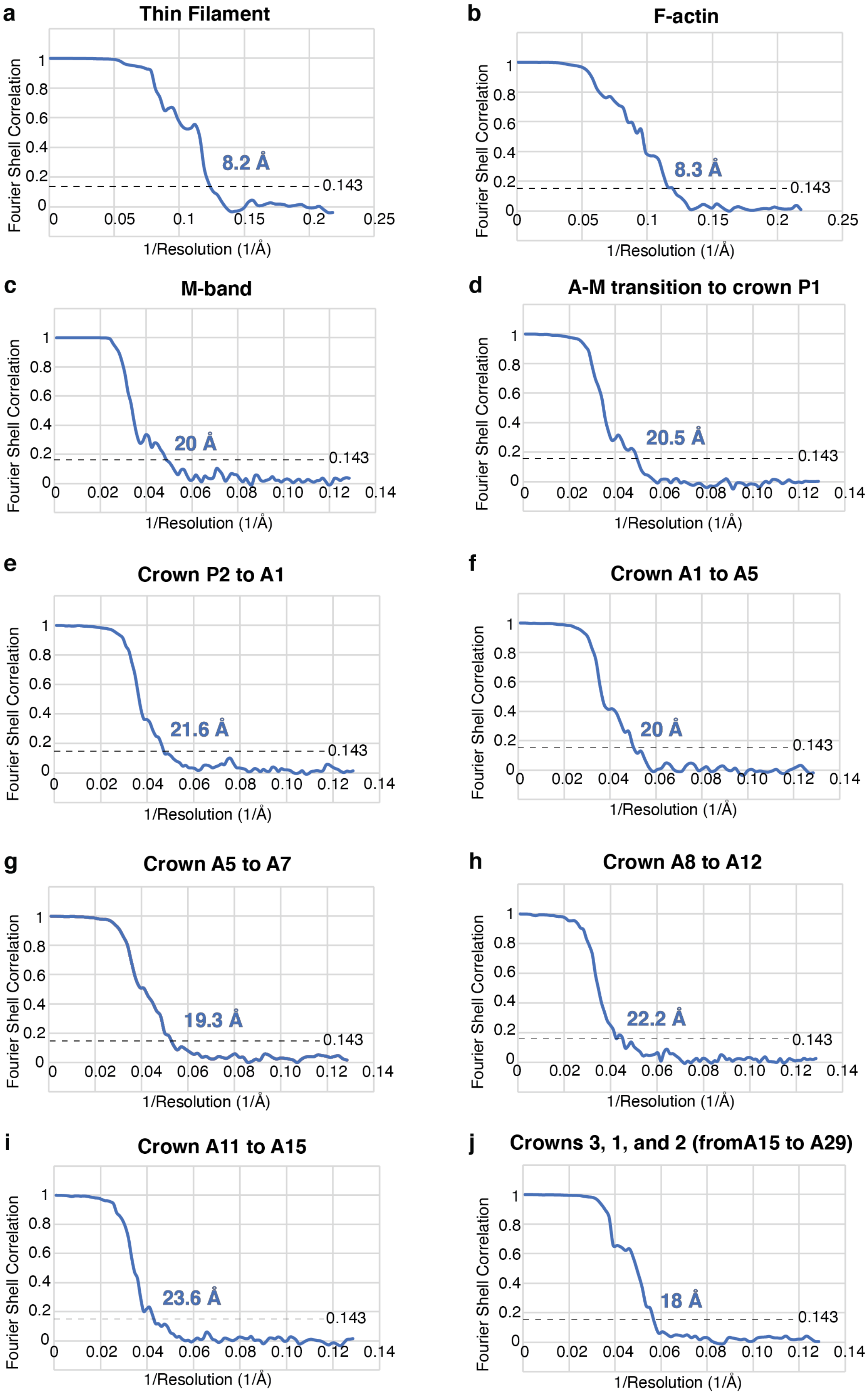

**Extended Data Figure 2** Gold-standard FSC curve of the cardiac native thin filament (**a**) and F-actin, after masking out tropomyosin (**b**)**.** Gold-standard FSC curves of the different segments of the relaxed thick filament: M-band (**c**)**,** A-M transition to crown P1 (**d**), from crown P2 to A1 (**e**), from crown A1 to A5 (**f**), from crown A5 to A7 (**g**), from crown A8 to A12 (**h**), from crown A11 to A15 (**i**), and the combined segments from crown A15 to A29 (**j**).

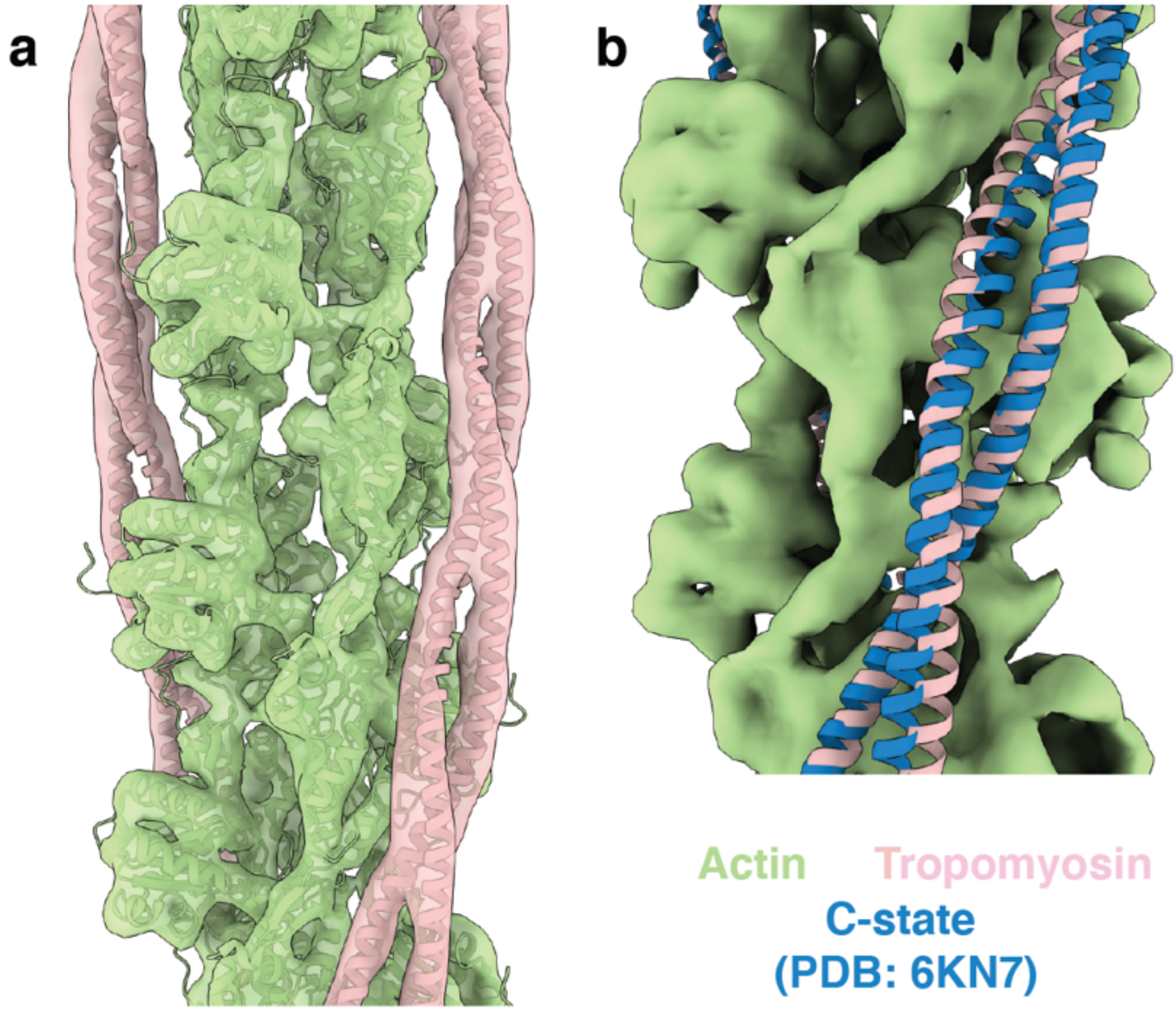


**Extended Data Figure 3 *In situ* structure of the thin filament in the $Ca^{2+}$ free state. a**, Subtomogram-averaged structure of the thin filament in the relaxed sarcomere. **b**, Model of the B-state of tropomyosin (PDB: 6KN7) is shown for comparison.

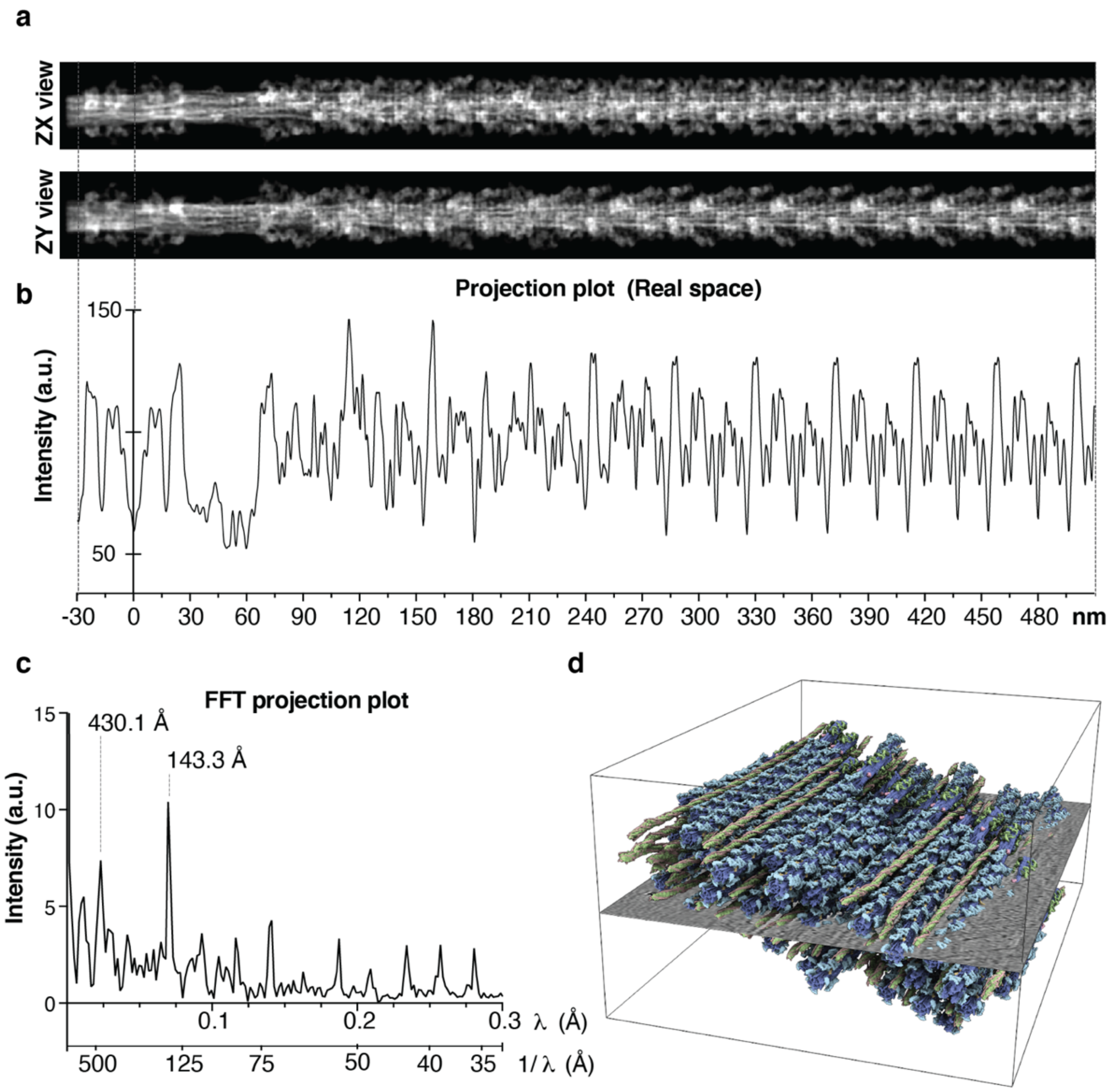


**Extended Data Figure 4 Thick filament density projection and profile. a**, The three-dimensional map of the thick filament was projected by summing the intensities of the voxels along the x axis. **b**, Intensity profile along the x axis, revealing the periodicity and spacing of the repeating units in the thick filament. **c**, The Fast Fourier Transform (FFT) of the intensity profile shows the peaks resulting from the repetitive nature of its component. The 430 Å peak is associated to the C-type SRs while the 143.3 Å peak is associated to the crowns layers and was used to calibrate the pixel size of the reconstruction. **d**, Tomographic volume of a cardiac sarcomere A-band showing thick and thin filaments back-plotted into the reconstructed volume with a single tomographic slice at the center for orientation.

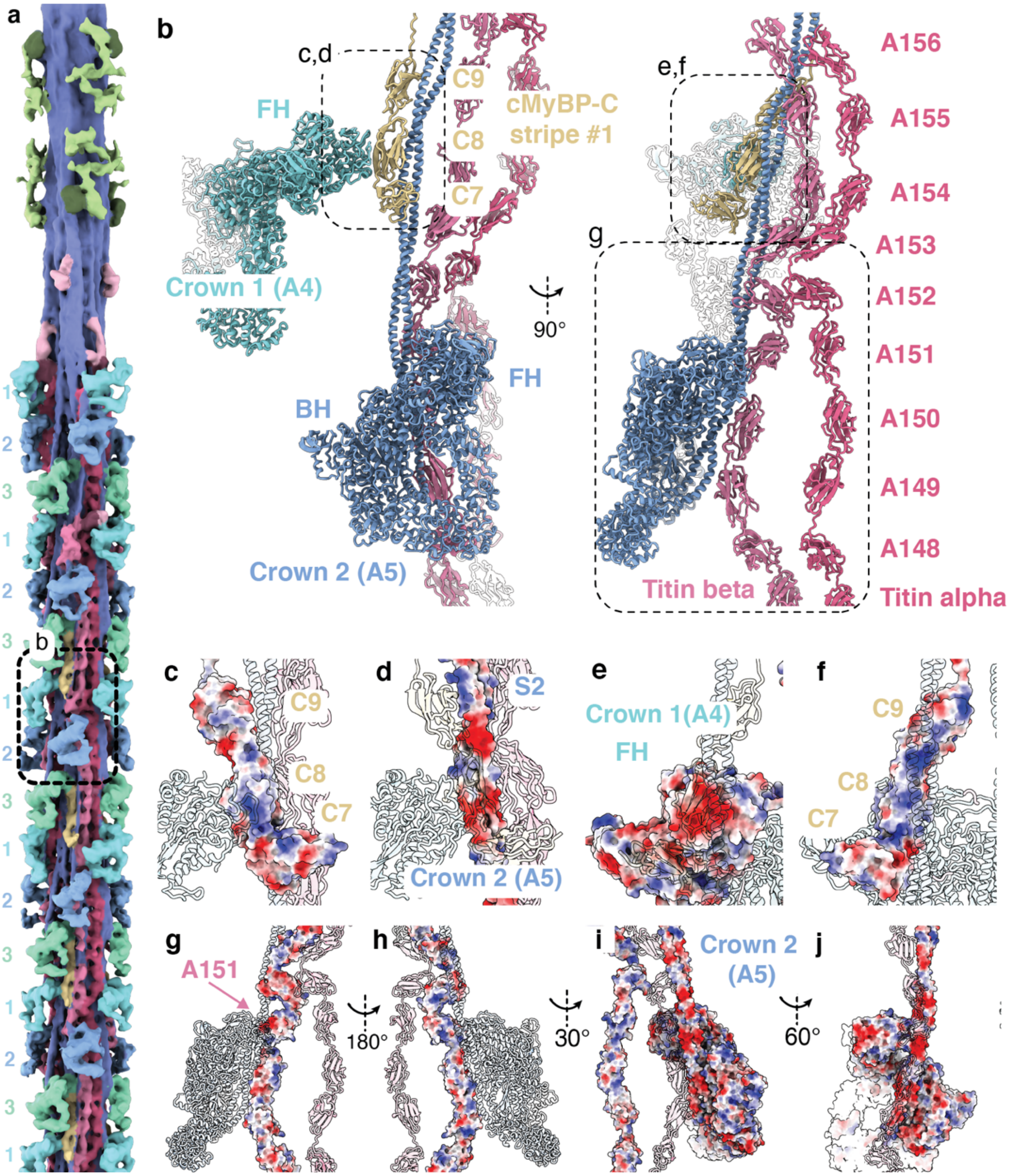


**Extended Data Figure 5 The OFF state of crowns 1 and 2 is stabilized by interactions with titin and cMyBP-C. a**, An overview of the thick filament from the M-band to the fourth cMyBP-C stripe is provided for context. **b**, Simplified molecular model of the titin CSR10. **c-f**, Crown 1 FH binds stably to the C8 domain of cMyBP-C via an extensive network of complementary charges (**c,d**), while C8 and C9 bind to the first 80 amino acids of crown 2 tails (**e,f**). **g-j**, The same tail region interact with titin beta domains (**g,h**). The OFF state of crown 2 is further stabilized by interactions with A151 (**i,j**).

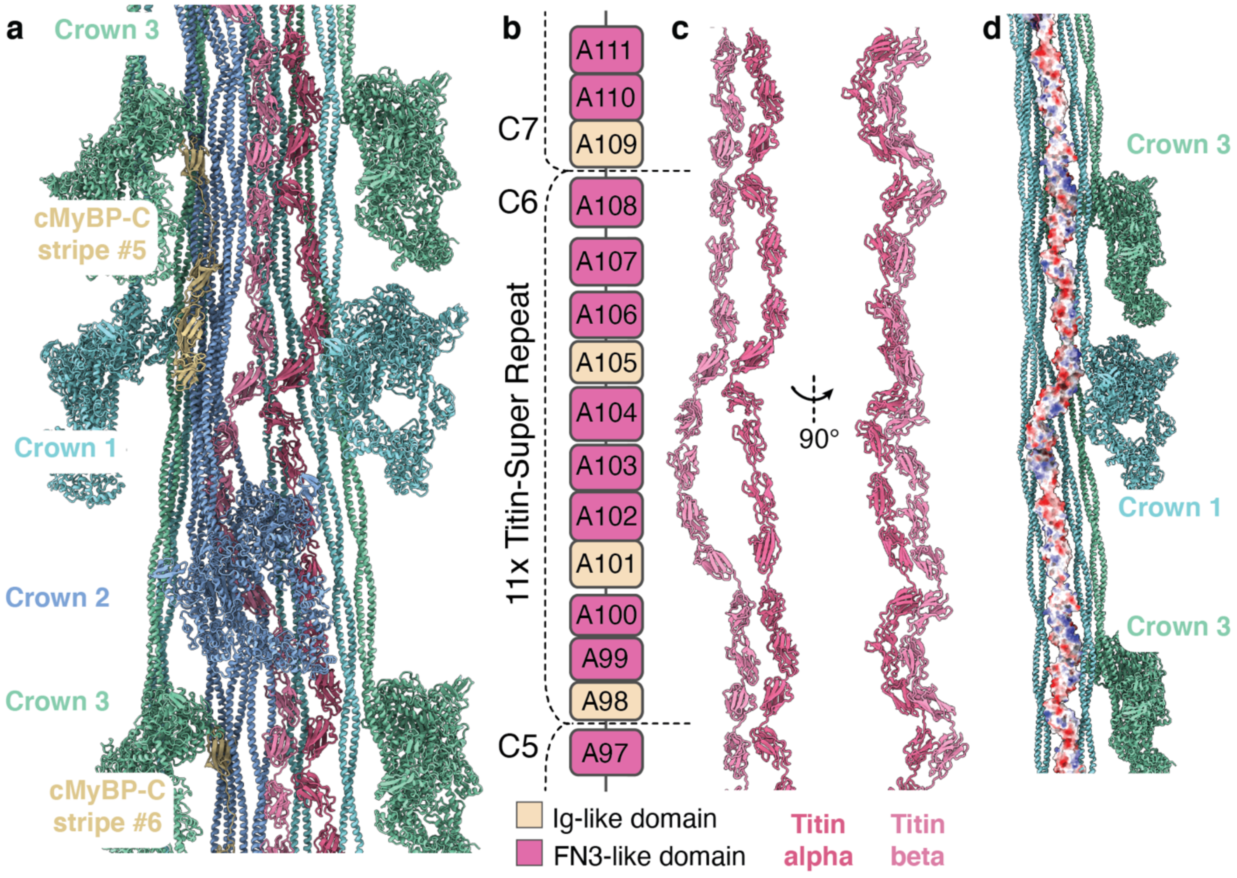

**Extended Data Figure 6 Titin organization in the center of the C-zone. a**, Simplified overview of a single asymmetrical unit at CSR6, showing the molecular arrangement of the myosin crowns, titin and cMyBP-C. **b**, Domain map of titin C-type super-repeat. **c**, Titin alpha and titin beta chain run parallel to each other and interact at the 1st, 3rd, 8th, and 11th CSR domains. **d**, Binding of titin to S2 and LMM of crowns 1 and 3 is reliant on a network of electrostatic interactions.

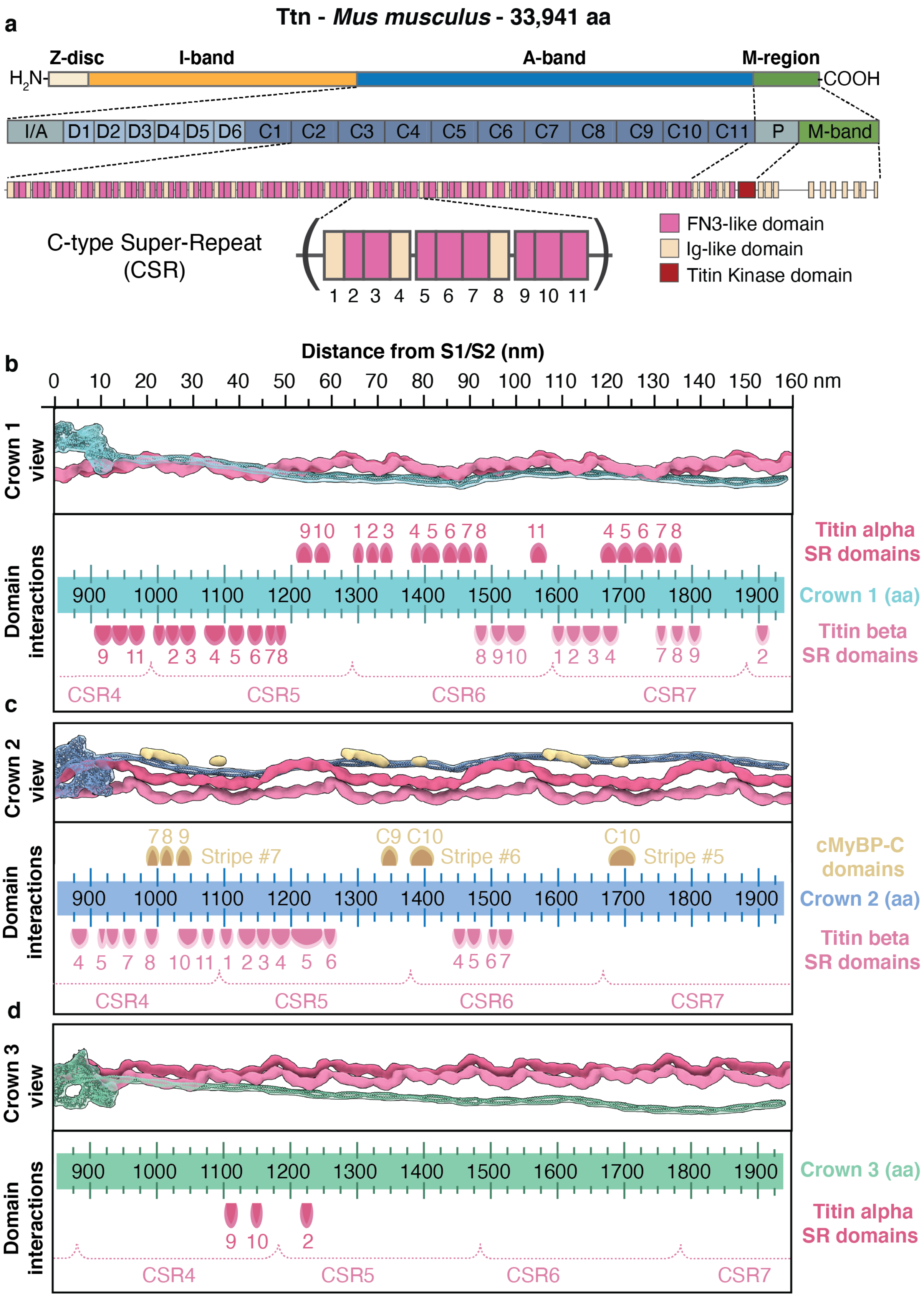
a
Ttn - Mus musculus - 33,941 aa
Z-disc
I-band
A-band
M-region
H2N-
-COOH
I/A D1 D2 D3 D4 D5 D6 C1 C2 C3 C4 C5 C6 C7 C8 C9 C10 C11 P M-band
C-type Super-Repeat (CSR)
1 2 3 4 5 6 7 8 9 10 11
FN3-like domain
Ig-like domain
Titin Kinase domain
b
Distance from S1/S2 (nm)
0 10 20 30 40 50 60 70 80 90 100 110 120 130 140 150 160 nm
Crown 1 view
Domain interactions
Titin alpha SR domains
Crown 1 (aa)
Titin beta SR domains
CSR4
CSR5
CSR6
CSR7
c
Crown 2 view
Stripe #7
Stripe #6
Stripe #5
cMyBP-C domains
Crown 2 (aa)
Titin beta SR domains
d
Crown 3 view
Crown 3 (aa)
Titin alpha SR domains

**Extended Data Figure 7 Titin's domain interactions with myosin tails. a,** Domain map of titin, showing its highly modular organization. The A-band comprises six D-type super-repeats in the distal zone (D-type SRs) and eleven 11x super-repeats in the central zone (C-type SRs). The proximal region (P-zone) contains the Titin Kinase domain (TK domain), while the M-band region presents ten Ig-like domains connected by seven disordered inter-spacing regions, m1-9 and is1-7, respectively. **b**, The model of crown 1 is shown alongside the densities of titin. To allow a clearer interpretation, a binding site map was generated to depict the amino acid residues of the myosin tail and their distance from the S1/S2 site. **c**, The arrangement and interactions of crown 2 are visualized alongside the densities of cMyBP-C and using the same approach as in (**b**). **d**, Similarly, the interaction of crown 3 with titin is shown.

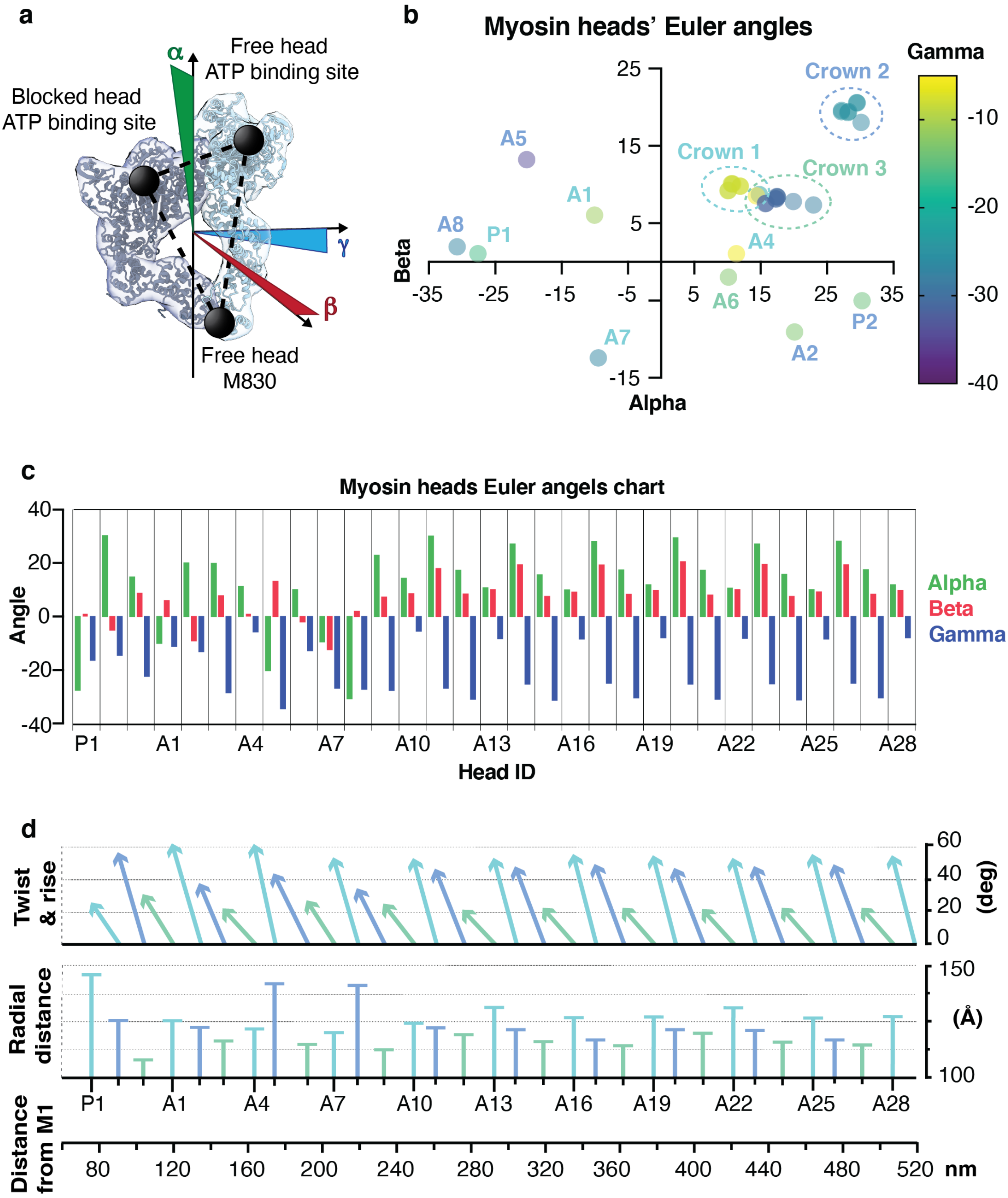


**Extended Data Figure 8 Position and orientation of myosin heads at each crown.** **a**, The arrangement of myosin heads deviates from a canonical helical symmetry. To quantitatively describe this arrangement, the thick filament was modeled as a cylinder, and the myosin IHMs were represented as triangles on its surface. The vertices of the triangle were determined by the coordinates of three points: the ATP-binding site in the FH, the same site in the BH, and the S1/S2 junction site. The Euler angles for the IHM are calculated relative to an equilateral triangle that has its topmost side perpendicular to the cylinder axis and parallel to its base. **b**, Colored scatter plot of the myosin crowns from P1 to A28. Alpha and beta angles are plotted along the x and y axes, respectively, while the gamma angle is encoded by the color gradient. Plotting of the IHMs generates three orientation clusters for the crowns from A10 to A28, while crowns associated to the P-zone or the first three C-type titin SR show more inconsistent orientation. **c**, A more detailed representation of (**b**) is shown in a colored histogram. **d**, Using the cylindrical coordinates for the triangles, we obtain the cylindrical coordinates for each myosin crown and plotted their twist, rise, and radial distance. Similar to the Euler angle distribution, a more consistent pseudo-helical pattern appears from A10 onwards. P1, A5, and A8, show outstanding orientation and radial distances, indicative of a more outward projected configuration.

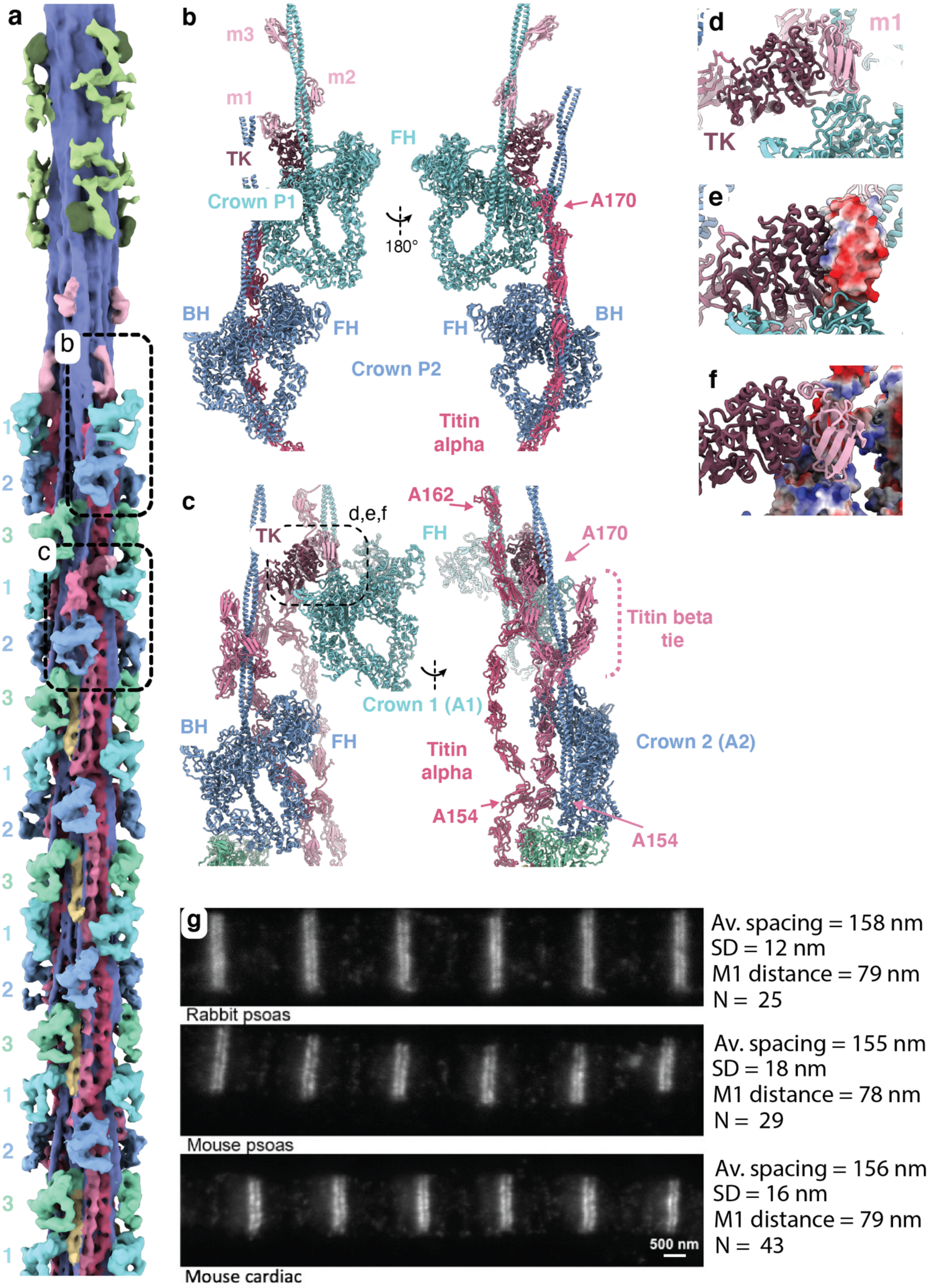


**Extended Data Figure 9 The P-zone of the thick filament. a**, An overview of the thick filament from the M-band to the fourth cMyBP-C stripe is provided for context. **b**, The P-zone of titin alpha is located behind the first two crowns, P1 and P2 while TK, m1, m2, and m3 are binding to P1's S1 and S2 domains. However, no density could be assigned to titin beta in this region. **c**, Titin beta chains clearly run alongside titin alpha chains until domain A154, and just behind the crown A2 motor domains. In this region, we observe additional globular densities, forming a flexible bow tie-shaped structure, which wraps around the tail of crown A2 and projects outward. These densities were assigned to titin beta P-zone domains. **d-f**, On top of crown A1, we observe an additional density that we attributed to the TK-m1 complex. In this configuration, m1 is positioned on top of the crown A2 BH and interacts with it mainly via electrostatic interactions. **g**, Super-resolution microscopy localisation of the TK domain showing STED imaging of titin kinase in myofibrils from different muscles labelled with affinity-purified anti-TK antibody. For all samples, we report the measurements of doublet spacing, placing TK at 79 nm from the M1 line. SD: standard deviation.

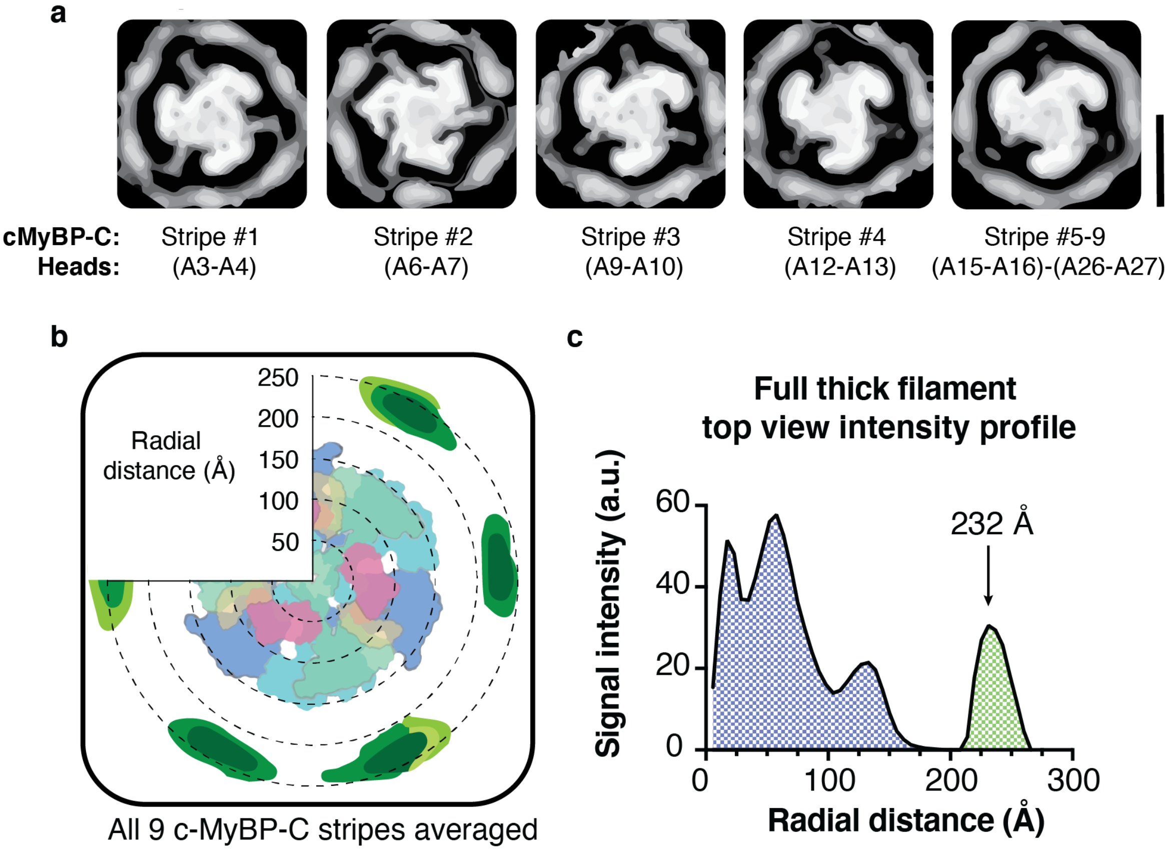


**Extended Data Figure 10 C-zone lattice organization. a**, The bridging domains of cMyBP-C exhibit a high degree of flexibility. To visualize the relative position of the thin filaments and the bridging cMyBP-C region with respect to the thick filament core, we centered each of the nine stripes individually on the cMyBP-C C7 plane and projected the resulting central 40 Å slab along the z-axis. Our analysis showed that stripes 4 to 9 projected into a region equidistant from both the negative azimuthal angle thin filament (NA-TF) and the positive azimuthal angle-thin filament (PA-TF), indicating that cMyBP-C may bind to either of them with equal likelihood. In contrast, stripes 2 showed a preferential orientation towards NA-TF, while stripe 1 exhibited a clear preferential bias towards PA-TF. **b,c,** We generated a single contour map by averaging all nine stripes together, which was then used to obtain a model for the determination of the average thin filament distance and azimuthal angles for the entire 430 Å repeat. Our analysis revealed that the center-to-center distance from thick to thin filament was 232 Å, as demonstrated by the radial profile of the signal intensity of a full 430 Å repeat. Scale bars: 25 nm.

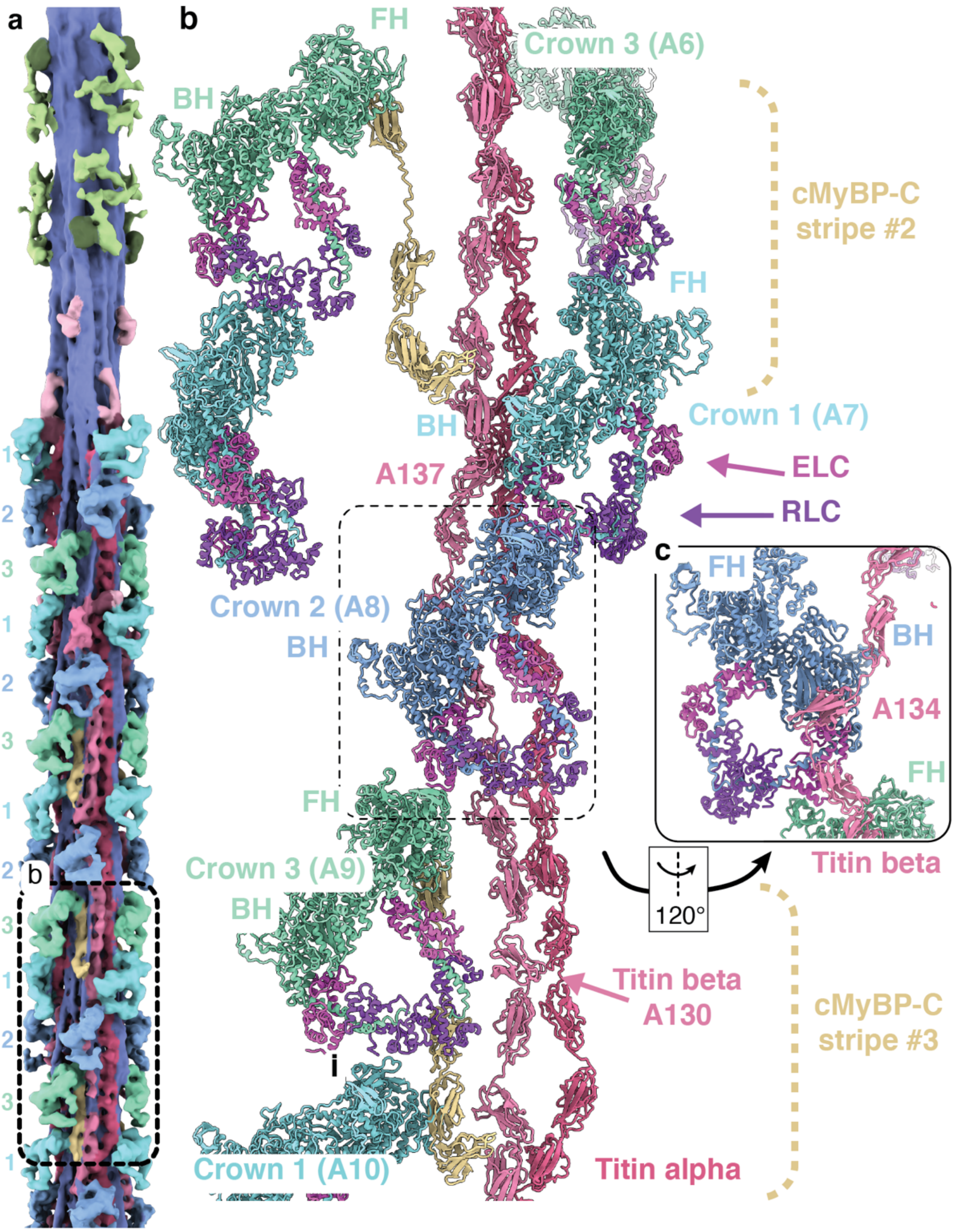


**Extended Data 11 Figure Myosin OFF state is stabilized by myosin light chains at CSR9.**

**a**, An overview of the thick filament from the M-band to the third cMyBP-C stripe is provided for context. **b**, Simplified overview of a single asymmetrical unit at CSR9, showing the molecular arrangement of the myosin crowns, titin and cMyBP-C. In contrast to all other cMyBP-C, at stripe #2 the crown 1 does not interact with cMyBP-C CTD, instead its FH binds to the RLC from crown 3 and its BH interacts with titin A137. Crown 2 orientation is also atypical as its FH interacts with the disordered region of Crown1's ELC and its BH binds to titin A134 (**c**). Similarly, crown 3 (A9)'s FH binds to the crown 2's ELC and it is also stabilized by an interaction of its RLC with titin beta A130.

**Extended Data Video 1 Tomogram of a sarcomere from the M-band to the C-zone.** The video shows the 3D organization of the relaxed cardiac sarcomere. The reconstructions of thick and thin filaments were back-projected into the tomogram to reveal their organization.

**Extended Data Video 2 Close-up of a segmented region in the tomogram showcasing the molecular details of the thick filament and its cMyBP-C links to the thin filament.**

**Extended Data Video 3 Tomogram of a C-zone in relaxed cardiac sarcomere.** cMyBP-C flexible links from thick to thin filament can be observed projecting from the thick filament at regular intervals of 430 Å.
